# Cytokinins induce prehaustoria coordinately with quinone and phenolic signals in the parasitic plant *Striga hermonthica*

**DOI:** 10.1101/2022.05.29.493927

**Authors:** Natsumi Aoki, Songkui Cui, Satoko Yoshida

**Author notes:** **Corresponding author:** Name: S. Yoshida, Address: Graduate School of Science and Technology, Nara Institute of Science and Technology, 8916-5 Takayama-cho, Ikoma, Nara, 630-0192 Japan.

## Abstract

Orobanchaceae parasitic plants are major threats to global food security, causing severe agricultural damage worldwide. Parasitic plants derive water and nutrients from their host plants through multicellular organs called haustoria. The formation of a prehaustorium, a primitive haustorial structure, is provoked by host-derived haustorium-inducing factors (HIFs). Quinones, including 2,6-dimethoxy-*p*-benzoquinone (DMBQ), and phenolics, including syringic acid, are of most potent HIFs for various species in Orobanchaceae, but except non-photosynthetic holoparasites, *Phelipanche* and *Orobanche* spp. On the other hand, cytokinin phytohormones was reported to induce prehaustoria in *Phelipanche ramosa*. However, little is known about whether cytokinins act as HIFs in the other parasitic species. Moreover, the signaling pathways for quinones, phenolics and cytokinins in prehaustorium induction are not well understood. This study showed that cytokinins act as HIFs in *Striga hermonthica* but not in *Phtheirospermum japonicum*. Using chemical inhibitors for each type of HIF, we demonstrated that cytokinins activate prehaustorium formation through a signaling pathway that overlaps with the quinone and phenolic HIF pathways in *S. hermonthica*. Host root exudates activated *S. hermonthica* cytokinin biosynthesis and signaling genes, and inhibitors blocking any of three types of HIFs perturbed the prehaustorium-inducing activity of exudates, indicating that host root exudates include a mixture of HIFs. Our study reveals the importance of cytokinins for prehaustorium formation in obligate parasitic plants.

## Introduction

Plants deprive other plants of nutrients are collectively called parasitic plants. Parasitic plants represent approximately 1% of angiosperms, including about 4,500 species, and the levels of host dependency and photosynthetic ability are varied among them (Yoshida et al. 2016). Those who cannot complete their life cycles without host plants in natural condition are called obligate parasites, whereas those can grow without host with their own photosynthetic activity are called facultative parasites. The obligate parasites can be further classified into hemi- and holo-parasites for those with or without photosynthesis ability, respectively, while all facultative parasites are photosynthetic hemiparasites. Parasitic plants in the Orobanchaceae family include all ranges of parasitic plants, from facultative parasites, obligate hemi- and holoparasites. Among them, the obligate hemiparasites *Striga*, and obligate holoparasites *Orobanche* and *Phelipanche*, are noxious weeds that cause significant reductions in crop yields (Mutuku et al. 2021). Development of efficient control methods for these weedy parasitic plants is of urgent issues, but fundamental knowledge of plant parasitism is still unraveled.

Host infection and nutrient acquisition from hosts by parasitic plants are achieved by a specialized invasive organ named the haustorium. All Orobanchaceae parasitic plants form haustoria on their roots. Haustorium initiation requires external haustorium-inducing factors (HIFs) produced by host plants (Goyet et al. 2019, Furuta et al. 2021). In response to HIFs, obligate hemi- and holoparasites, such as *Striga* and *Phelipanche*, terminate radicle elongation and induce radial cell expansion and cell division to form a swollen-tip structure, termed a terminal prehaustorium (Fernández-Aparicio et al. 2016, Cui et al. 2018, Furuta et al. 2021). Facultative hemiparasites that do not rely on a host to complete their life cycle, including *Triphysaria versicolor* and *Phtheirospermum japonicum*, maintain root apical meristems even after HIF recognition and induce prehaustoria in the upper root transition zone, resulting in multiple lateral prehaustoria along a root (Yoshida et al. 2016). Prehaustoria further invade host roots and develop xylem bridges between the host and parasite, leading to vascular connections for material transfer via the development of mature haustorium structures (Masumoto et al. 2021).

Several HIFs and their structure-activity specificities have been reported in Orobanchaceae members (Goyet et al. 2019). A quinone, 2,6-dimethoxy-*p*-benzoquinone (DMBQ), is a well-known HIF that can induce prehaustoria of many Orobanchaceae hemiparasites, including the obligate hemiparasite *Striga* spp. and facultative hemiparasites *Triphysaria* and *Phtheirospermum*. In addition, phenolic compounds with structural similarity to lignin monomers, including syringic acid and acetosyringone, have prehaustorium induction activity in these species with similar or lower activity than DMBQ (Cui et al. 2018). These phenolic HIFs commonly have a hydroxyl group at position 4 of a benzene ring and methoxy groups at positions 3 and/or 5. It appears that the number of methoxy groups affects the prehaustorium-induction activity in *S. hermonthica* and *P. japonicum* (Cui et al. 2018). Notably, quinone or phenolic HIFs are not highly active HIFs in obligate holoparasitic *Phelipanche* species (Goyet et al. 2017, Westwood et al. 2010). DMBQ can induce prehaustoria in *Phelipanche ramosa* but only at a concentration higher than 500 µM (Fernandez-Aparicio et al. 2021). Instead, cytokinin phytohormones, structurally unrelated to quinones or phenolics, reportedly induce early haustorial structures in *P. ramosa*. Furthermore, their *Brassica napus* host plants exude cytokinin-like allelochemicals with haustorium-inducing activity (Goyet et al. 2017). Other studies have reported that cytokinins induce prehaustorium-like structures in *Striga asiatica* and *Triphysaria vesicolor* (Keyes et al. 2000, Wrobel and Yoder, 2001); however, no additional information has been reported to date. Nevertheless, these observations have raised the possibility that cytokinins may be common HIFs for both hemiparasites and holoparasites.

Genes mediating quinone-type HIF signals were reported in several Orobanchaceae species. Genes encoding quinone oxidoreductase (QR) and the general transcription factor PIRIN were upregulated after application of DMBQ or host root exudates to *T. versicolor* and *P. japonicum* (Bandaranayake et al. 2010, 2012, Ishida et al. 2017). In *T. versicolor*, QR1 catalyzes NADPH-dependent quinone reduction, and this process was proposed to be important for transducing quinone signals for prehaustorium induction (Bandaranayake et al. 2010, Wrobel et al. 2002). Knockdown of QR1 or PIRIN resulted in reduced prehaustorium induction in *T. versicolor* (Bandaranayake et al. 2010, 2012). The auxin biosynthesis gene, *YUCCA3*, is expressed at the prehaustorium formation site, providing the necessary auxin required for prehaustorium formation in *P. japonicum* (Ishida et al. 2016). Expansin genes are upregulated by DMBQ treatment in *S. asiatica* (O’Malley and Lynn 2000). Moreover, the DMBQ-sensing genes were recently identified. DMBQ is recognized *via* a receptor-like kinase, CANNOT RESPOND TO DMBQ 1 (CARD1) in *A. thaliana*, leading to an increase in intercellular [Ca^2+^] and regulation of stomatal closure and immunity (Laohavisit et al. 2020). *P. japonicum* and *S. asiatica* have three CARD1 homologues that can complement [Ca^2+^] elevation defects in Arabidopsis *card1*, indicating functional conservation of parasitic plant CARD1 in DMBQ perception (Laohavisit et al. 2020). Also, inhibitor assays suggest that [Ca^2+^] regulation is important for DMBQ-induced prehaustorium formation in *P. japonicum* (Laohavisit et al. 2020). Whether CARD1 homologs are present and functional in obligate holoparasites *Phelipanche* and *Orobanche spp*. remains unclarified.

Phenolic HIFs have been proposed to act as DMBQ precursors during prehaustorium induction, as syringic acid or sinapic acid are oxidatively converted to DMBQ *in vitro* by peroxidases, presumably produced by parasitic plants (Kim et al. 1998). It remains possible that phenolic HIFs may act as signaling molecules in prehaustorium induction. The gram-negative bacterium *Rhizobium radiobacter* (syn. *Agrobacterium tumefaciens*) recognizes acetosyringone produced from wounded plant cell walls *via* the *virA* sensor to induce downstream *Vir* gene expression, enabling the transfer of DNA fragments into plant genomes (Gelvin, 2000). Although direct binding of acetosyringone and VirA was not confirmed, the acetosyringone analog α-bromoacetosyringone (ASBr) functions as a competitive inhibitor to suppress *Vir* gene expression (Hess et al. 1991). Interestingly, acetosyringone induced prehaustoria in *P. japonicum* and *S. hermonthica* (Cui et al. 2018). Whether these phenolic compounds are sensed by direct sensor(s) or are converted to quinones in parasitic plants remains questionable.

Cytokinins are well-known plant growth regulators. In plant roots, cytokinins promote vascular differentiation and nodulation, regulate nutrient uptake and inhibit lateral root formation and root elongation (Kieber and Schaller 2018, Werner and Schmülling 2009). Cytokinin signals are transduced *via* a two-component system. In *Arabidopsis thaliana*, ARABIDOPSIS HISTIDINE KINASE 2 (AHK2), AHK3 and AHK4 were identified as cytokinin receptors (Inoue et al. 2001, Nishimura et al. 2004). These histidine kinases have cytokinin-binding cyclase/Histidine kinase-associated sensing extracellular (CHASE) domains (Kakimoto 2003). Binding to cytokinin activates phosphorylation of histidine in the kinase domain, and the phosphorylation signal is transferred to histidine on ARABIDOPSIS HIS PHOSPHOTRANSFER PROTEINS (AHPs), which then transfers the phosphoryl group to the receiver domains of ARABIDOPSIS RESPONSE REGULATORS (ARR) (To and Kieber 2008, Ferreira and Kieber 2005).

Orobanchaceae parasitic plants can respond to various types of molecules as HIFs, yet the signaling pathways for each HIF and their relationships are largely unexplored. Moreover, although DMBQ was isolated from *Sorghum* extracts (Chang and Lynn 1986), DMBQ is barely detected in *Arabidopsis thaliana* root exudates, which have high prehaustorium induction activities (Wang et al. 2020). Therefore, the actual molecules in host root exudates responsible for prehaustorium induction are still unknown. In this study, we show that cytokinins function in prehaustorium induction in *S. hermonthica* but not *P. japonicum*. Taking advantage of the available inhibitors for quinone, phenolics and cytokinins, we evaluated the relationship of their signaling pathways in prehaustorium formation and demonstrated the presence of potential HIFs in host root exudates.

## Results

### Cytokinins induce prehaustoria in Striga hermonthica

Previous studies showed that cytokinins induce the formation of prehaustoria (early haustorial structures) in *P. ramosa* and prehaustorium-like structures in *S. asiatica* (Goyet et al. 2017, Keyes et al. 2000), implying that cytokinins might be common HIFs in Orobanchaceae. Therefore, we tested the prehaustorium-inducing activity of five natural or synthetic cytokinins, including 6-benzylaminopurine (BA), kinetin (KIN), N6-isopentenyladenine (2iP), and *trans*-zeatin (tZ), and a synthetic derivative thidiazuron (TDZ), on the obligate hemiparasite *S. hermonthica* and the facultative hemiparasite *P. japonicum*. We tested a range of cytokinin concentrations on germinated *S. hermonthica* seedlings and measured prehaustorium formation (Figs. 1 and 2). For comparison, previously identified HIFs, including DMBQ, syringic acid (SyA), acetosyringone (AS) and host root exudates from rice or *Arabidopsis*, were tested using the same conditions. All tested cytokinins induced root tip swelling and proliferation of haustorial hairs on the surface of swelling sites, a typical prehaustorial structure observed after treatment with previously known HIFs or host root exudates (Fig. 1B). Dose-response experiments showed that all tested cytokinins had higher prehaustorium-inducing activities than DMBQ, AS or SyA, of which TDZ had the highest and KIN had the lowest activities (Fig. 2). Prehaustorium formation stimulated by DMBQ, AS and SyA reached about 80% at 1 µM, 20 µM and 5 µM, respectively (Fig. 2A), whereas similar levels were achieved by nanomolar ranges of cytokinins (10 nM, 30 nM, 80 nM, 250 nM, 400 nM by TDZ, tZ, 2iP, BA and KIN, respectively; Fig. 2B).

**Fig. 1.**
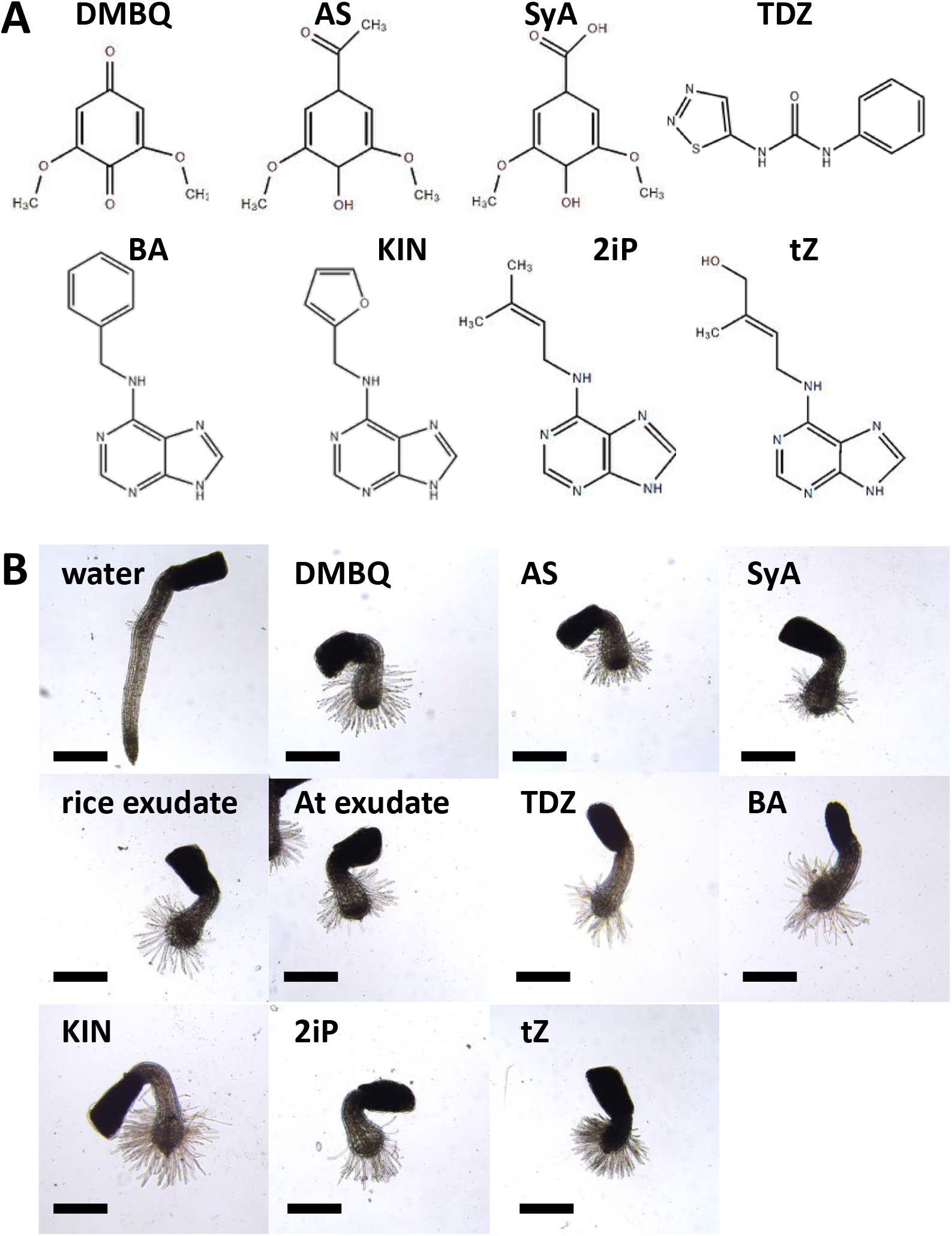
Prehaustorium-inducing activity of several HIFs including quinone-type, phenolic-type, and cytokinin-type HIFs. A, The chemical structures of HIFs tested in this study. B, *S. hermonthica* prehaustoria induced by rice root exudate, *A. thaliana* (At) root exudate, DMBQ (10 μM), AS (50 μM), SyA (20 μM), TDZ (200 nM), BA (500 nM), KIN (500 nM), 2iP (200 nM) and tZ (200 nM). Scale bars=500 μm.

**Fig. 2.**
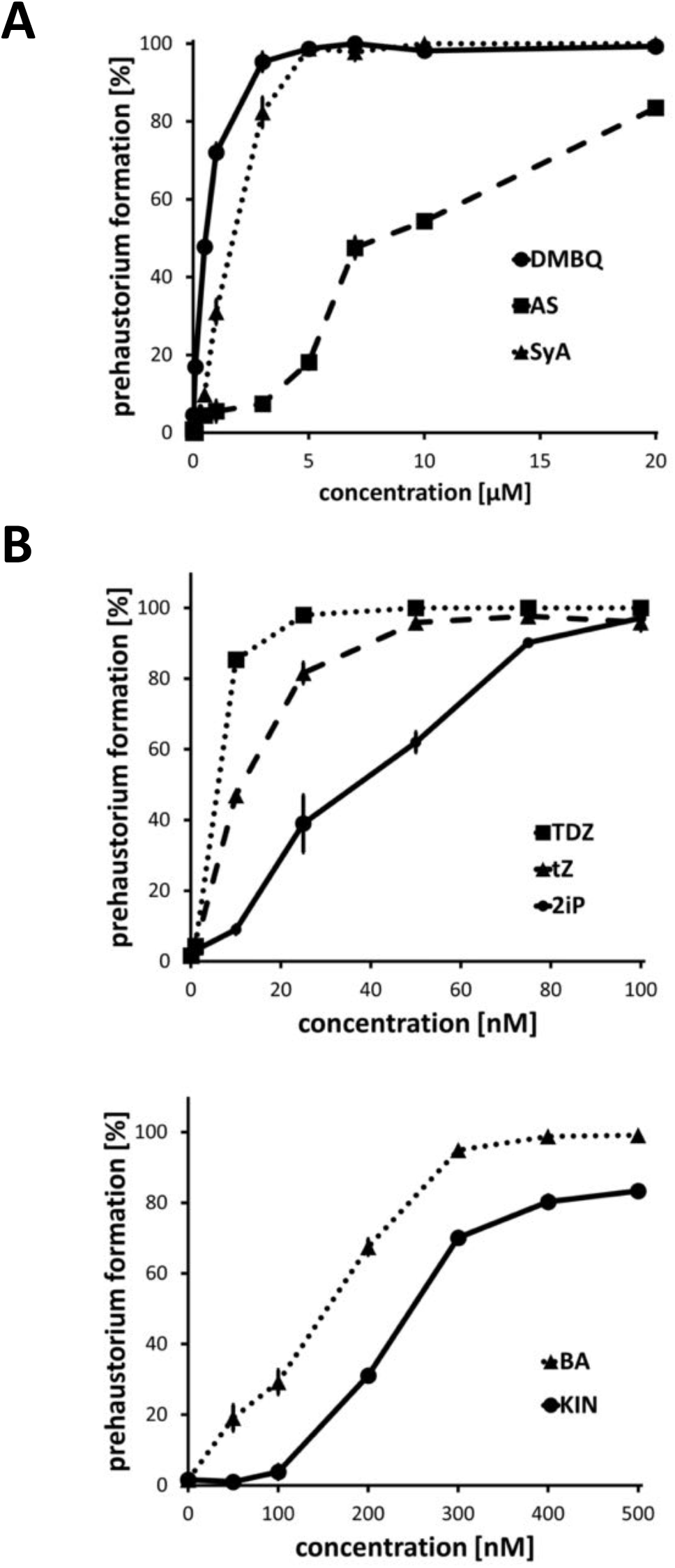
Concentration-dependent prehaustorium-inducing activity of HIFs. A, Prehaustorium-inducing activity of DMBQ, AS and SyA. B, Prehaustroium-inducing activity of cytokinins (BA, KIN, tZ, TDZ and 2iP). *S. hermonthica* seedlings were treated with HIFs for 24 h. Data represent the mean ± standard error (n=3).

### Cytokinins do not induce prehaustoria in Phtheirospermum japonicum

We next tested the effects of cytokinins on the facultative parasite *P. japonicum*, but none of the tested cytokinins induced prehaustoria after 7 days of treatment at 500 nM (Figs. S1A and S1B). To confirm that *P. japonicum* can respond to cytokinins, we observed *P. japonicum* morphological changes after cytokinin treatments. As was the case for Arabidopsis (Smets et al. 2005, Werner et al. 2001), cytokinins altered root morphology in *P. japonicum* (Figs. S1). TDZ suppressed lateral root formation by one-half compared to that of the water treatment (Fig. S1C), and *t*Z, TDZ and 2iP significantly inhibited primary root growth (Fig. S1D). Moreover, BA or KIN treatments increased root hair proliferation, whereas tZ or TDZ induced root greening (Fig. S1B). These observations indicate that *P. japonicum* can respond to exogenously applied cytokinins as growth regulators but not as HIFs.

### Dissection of signaling pathways for prehaustorium induction in S. hermonthica

The above-described experiments indicate that *S. hermonthica* can induce prehaustoria in response to three types of HIFs: quinones such as DMBQ, phenolics such as AS and syringic acid, and cytokinins such as tZ and TDZ. However, how each type of HIF signals is transduced and interacts with each other in inducing prehaustorium are unknown. To understand the relationships of haustorium induction signaling pathways activated by each type of HIF, we investigated the effect of chemical inhibitors on *S. hermonthica* prehaustorium formation. Tetrafluoro-1,4-benzoquinone (TFBQ) (Fig. 3A), a DMBQ analog, has been used as an inhibitor for DMBQ-induced prehaustorium formation in *S. asiatica, T. versicolor* and *P. japonicum* (Smith et al. 1996, Wang et al. 2019, Laohavisit et al. 2020). We found that TFBQ not only inhibits DMBQ-induced prehaustorium formation but also AS-, SyA-or cytokinin-induced prehaustorium formation (Figs. 3D, S2A, S3A-C). Prehaustorium formation was gradually restored by increasing the concentration of DMBQ, AS or *t*Z, indicating that TFBQ perturbs the activity of these HIFs in a competitive manner (Fig. 3D). ASBr (Fig. 3B) is an analog of AS (Fig. 1A) and was reported to inhibit AS-dependent induction of *Vir* gene expression in *Agrobacterium* (Hess et al. 1991). Like TFBQ, the application of ASBr strongly inhibited prehaustorium formation by all the HIFs, DMBQ, AS, SyA and cytokinins (Figs. 3E, S2B and S3A-C). As cytokinin inhibitors, we tested two cytokinin analogs, LGR-991 (Fig. 3C) and PI-55 (Fig. S4A), that block cytokinin binding to AHKs, the cytokinin receptors in *Arabidopsis*, in a cytokinin-competitive manner (Nisler et al. 2010, Spíchal et al. 2009). Although PI-55 was reported to suppress prehaustorium formation in *P. ramosa* (Goyet et al. 2017), we found that PI-55 or LGR-991 induced prehaustoria in *S. hermonthica* (Fig. S4), most likely due to their weak cytokinin agonist activity (Spíchal et al. 2009). Nevertheless, we were able to use 10 µM LGR-991 to test the effects of a cytokinin inhibitor on prehaustorium induction because LGR-991 provoked prehaustorium formation only at concentrations higher than 20 µM (Fig. S4B). LGR-991 at 10 µM inhibited prehaustorium formation induced by cytokinins (Figs. 3F and S3C), suggesting that cytokinin perception by AHK homologs are necessary for prehaustorium induction by cytokinins in *S. hermonthica*. Interestingly, LGR-991 also inhibited AS- and SyA-induced prehaustoria (Figs. 3F, S2C and S3A-B), although the inhibitory effects were lower than those to cytokinins (Figs. 3D and 3E). However, LGR-991 did not affect prehaustoria formation induced by DMBQ (Figs. 3F and S2C), indicating that cytokinin perception *per se* is dispensable in DMBQ-induced prehaustorium formation in *S. hermonthica*. These findings suggest that cytokinin and phenolic pathways overlap at least partially in prehaustorium induction.

**Fig. 3.**
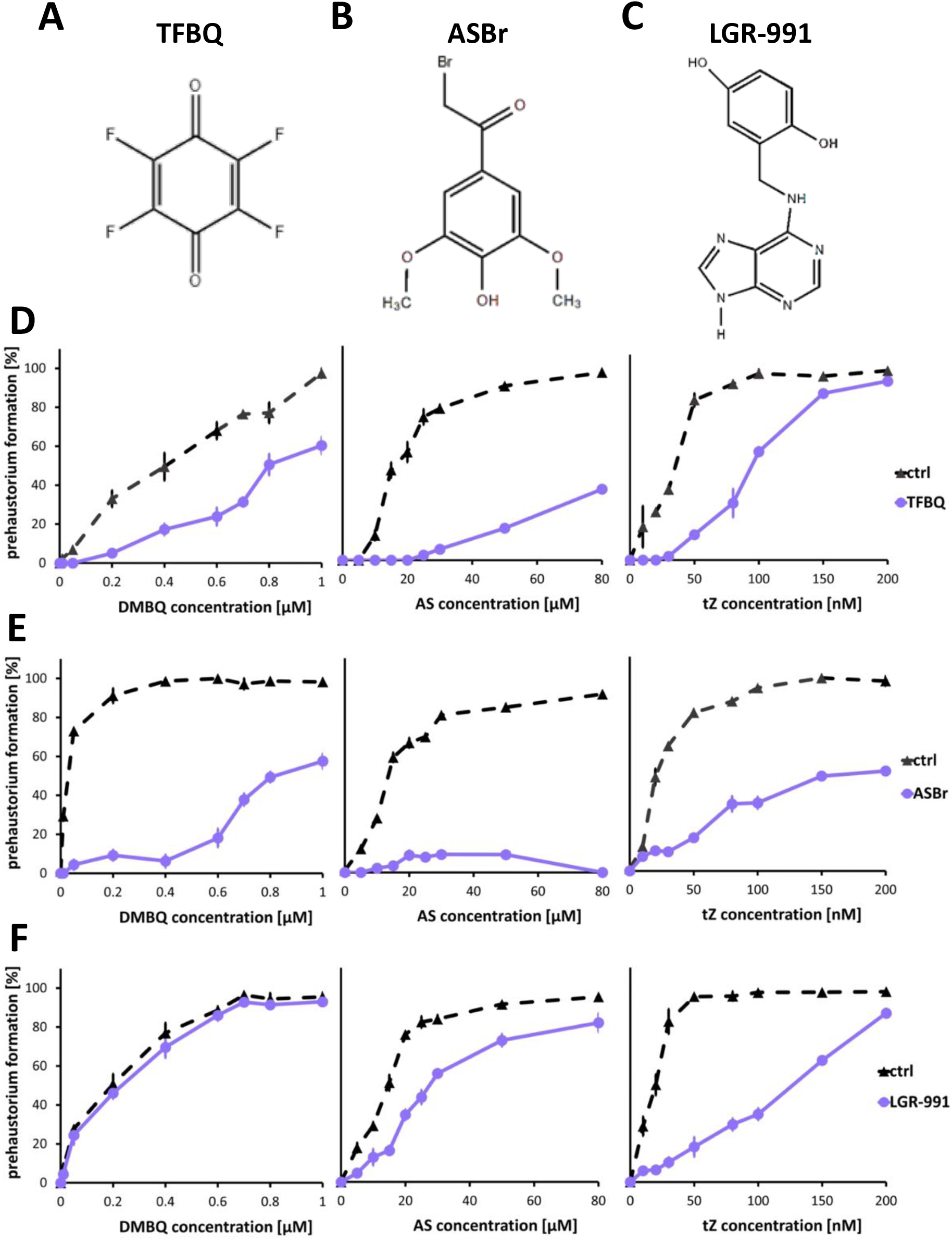
Inhibitory effects on *S. hermonthica* prehaustorium induction by DMBQ, AS and tZ. A-C, The chemical structures of inhibitors: TFBQ, a DMBQ analog; ASBr, an AS analog; LGR-991, a cytokinin inhibitor. D-F, The inhibitory effect of 20 µM TFBQ (D), 10 µM ASBr (E) and 10 µM LGR-991 (F) on DMBQ, AS or tZ. *S. hermonthica* was treated with HIFs and inhibitors for 24 h. Data represent the mean ± standard error (n=3).

Because TFBQ and ASBr inhibit prehaustorium formation by cytokinins, a question was raised as to whether these inhibitors inhibit general cytokinin-signaling. Thus, we tested the effects of TFBQ and ASBr on cytokinin-induced root growth suppression and promotion of hypocotyl elongation. Because *S. hermonthica* does not form roots or hypocotyls without host infection, we used *P. japonicum* for this analysis. *t*Z treatment suppressed primary root length ad lateral root numbers, and increased hypocotyl length in *P. japonicum* (Fig. S5A-C). These growth alterations by *t*Z were attenuated by LGR-991 but not by TFBQ or ASBr (Fig. S5A-C). These observations imply that TFBQ and ASBr inhibit cytokinin-induced prehaustorium formation but not cytokinin-mediated growth regulation in parasitic plants.

We also tested the effects of three inhibitors on *P. japonicum* prehaustorium formation. Prehaustorium number after DMBQ treatment was significantly reduced by treatment of either of TFBQ, ASBr or LGR-991 (Fig. S5D). Similarly, all inhibitors perturbed prehaustorium formation by AS treatment (Fig. S5D). Although cytokinins are not prehaustorium inducers in *P. japonicum* (Fig. S1), LGR-991 still showed strong inhibitory effects, indicating the importance of cytokinin signaling for prehaustorium formation in *P. japonicum*.

### Gene expression downstream of each HIF

Inhibitor assays indicated that the signaling pathways of cytokinins overlap with other HIFs at downstream, but the precise relationships of these HIF signaling are still unclear. To dissect the relationship of HIF signaling, we investigated gene expression in their response to three types of HIFs. We used RT-qPCR to measure the expression of four genes, QR2, PIRIN, YUCCA3 and EXPB1, which were previously reported to be DMBQ-inducible genes in Orobanchaceae parasitic plants (Bandaranayake et al. 2010, 2012, Ishida et al. 2016, 2017, O’Malley and Lynn 2000). In addition, we analyzed the expression of chitinases (chitinase and a class iv chitinase), the plant defense-related genes (Sharma et al. 2011) reported to be highly expressed at the early stages during host infection and by DMBQ treatment in *S. hermonthica* (Yoshida et al. 2019). *ShQR2* was upregulated 6 h or 24 h after DMBQ or SyA treatment, respectively (Fig. 4A). Similarly, *ShPIRIN* was upregulated by more than 2-fold at 1 and 6 h after DMBQ treatment and at 24 h after SyA treatment. On the other hand, tZ treatment induced expression of *ShPIRIN* but not *ShQR2* (Fig. 4A). *ShYUC3* expression increased at 6 h by all HIFs, including tZ, and a higher expression level was maintained only by SyA at 24 h (Fig. 4A). The expression of *ShEXPB1* was significantly upregulated by SyA and slightly by DMBQ, AS at 24 h but not by tZ (Fig. 4A). The chitinase homologue was upregulated by DMBQ at 6 h and by SyA at 24 h, whereas it was slightly upregulated by tZ at 6h without statistical significance (Fig. 4A). The class iv chitinase gene expression was upregulated by DMBQ, AS and SyA at 24 h but not by tZ. These results indicate that cytokinins induce prehaustorium formation by activating a subset of genes downstream of quinones and phenolics. Thus, cytokinins are not placed upstream of DMBQ or phenolics.

**Fig. 4.**
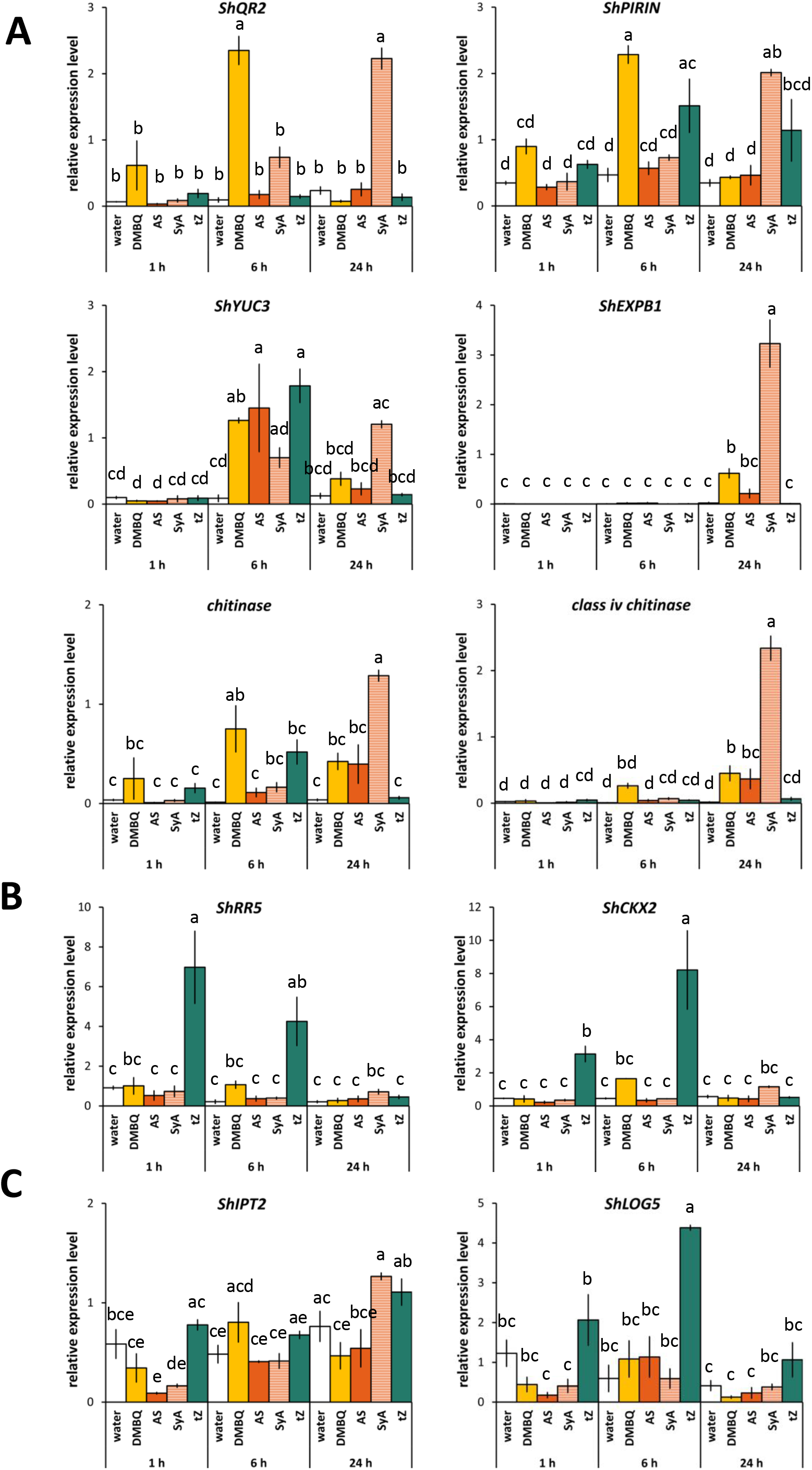
Expression of genes involved in prehaustorium formation, cytokinin signaling and cytokinin biosynthesis in response to several HIFs. A, Expression of DMBQ marker genes (*ShQR2, ShPIRIN, ShYUC3, ShEXPB1*, chitinase and class iv chitinase). B, Expression of genes involved in cytokinin signaling (*ShRR5* and *ShCKX2*). C, Expression of genes involved in cytokinin biosynthesis (*ShLOG5* and *ShIPT2*). *S. hermonthica* was treated with DMBQ (10 µM), AS (50 μM), SyA (20 μM) or tZ (200 nM) for 24 h. Data represent the mean ± standard error (n=3). Letters indicate statistically significant differences (Tukey’s test, *p*<0.05).

Next we tested the possibility that cytokinin signaling can be placed downstream of quinone or phenolic HIF signaling pathways. The expression of *S. hermonthica* genes homologous to the cytokinin-responsive *ARR5* and *CKX2* in *A. thaliana* (Bhargava et al. 2013) were investigated by RT-qPCR. As expected, expression of *ShARR5* and *ShCKX2* was significantly increased at the early time points (1 and 6 h post-treatment) by tZ (Fig. 4B). On the other hand, neither DMBQ, SyA nor AS significantly affected the expression of *ShRR5* and *ShCKX2* (Fig. 4B). These findings imply that quinone and phenolics induce prehaustorium formation without activating the canonical cytokinin signaling pathway.

To investigate whether quinones or phenolics affect cytokinin biosynthesis, we measured the expression of cytokinin biosynthesis genes in *S. hermonthica* in response to DMBQ, SyA or AS (Fig. 4C). Adenosine phosphate-isopentenyltransferases (IPTs) and cytokinin riboside 5’-monophosphate phosphoribohydrolases encoded by *LONELY GUY (LOG)* are known as key genes in the cytokinin biosynthesis pathway (Kamada-Nobusada and Sakakibara 2009). We searched for homologous genes of IPTs and LOGs in the *S. hermonthica* transcriptome (Yoshida et al., 2019) and found 9 and 10 homologues, respectively (Fig. S6). Because *AtLOG5* and *AtIPT2* encode functional enzymes responsible for major cytokinin biosynthesis activity in root apical meristems and vascular tissues (Miyawaki et al. 2004, Kuroha et al. 2009), we chose their orthologues, *ShLOG5* and *ShIPT2*, for expression analyses. *ShIPT2* expression was induced by neither of HIFs (Fig. 4C). *ShLOG5* was upregulated by tZ but not by DMBQ, AS and SyA (Fig. 4C). Our results further imply that quinone and phenolic HIFs induce prehaustorium formation without activating cytokinin biosynthesis and signaling.

### HIFs in host root exudates

In natural conditions, *S. hermonthica* initiates haustorium formation in response to nearby host roots. Therefore, host root exudates must contain HIFs, but the identities of host-derived HIFs are unclear. To answer this question, we took advantage of available inhibitors and investigated the effects of TFBQ, ASBr and LGR-991 on prehaustorium formation stimulated by host root exudates (Fig. 5A). We choose two plant species, rice and Arabidopsis; rice as a representative of natural hosts for *S. hermonthica* and Arabidopsis as a plant having prehaustorium inducing activity for *S. hermonthica* although it is non-host in natural condition (Yoshida and Shirasu, 2009). All inhibitors reduced prehaustorium formation in treatments of rice or Arabidopsis root exudates. Whereas rice or Arabidopsis root exudates led to prehaustorium induction levels of more than 80%, the addition of TFBQ reduced prehaustoria by less than 10% in both (Figs. 5A and S2D). The inhibitory effects of ASBr were different between rice and Arabidopsis exudates; ASBr had a stronger effect on rice exudates with an induction level of less than 10% compared with Arabidopsis exudate (40-60%) (Figs. 5A and S2E). Treatment of LGR-991 decreased the prehaustorium formation rates by either exudate to 40–60% (Figs. 5A and S2F). Because LGR-991 showed little inhibition of DMBQ-induced prehaustoria formation in *S. hermonthica* (Fig. 3F), these results indicate that root exudates of both rice and Arabidopsis contain not solely quinone-type HIFs but may also contain phenolic or cytokinin HIFs or both.

**Fig. 5.**
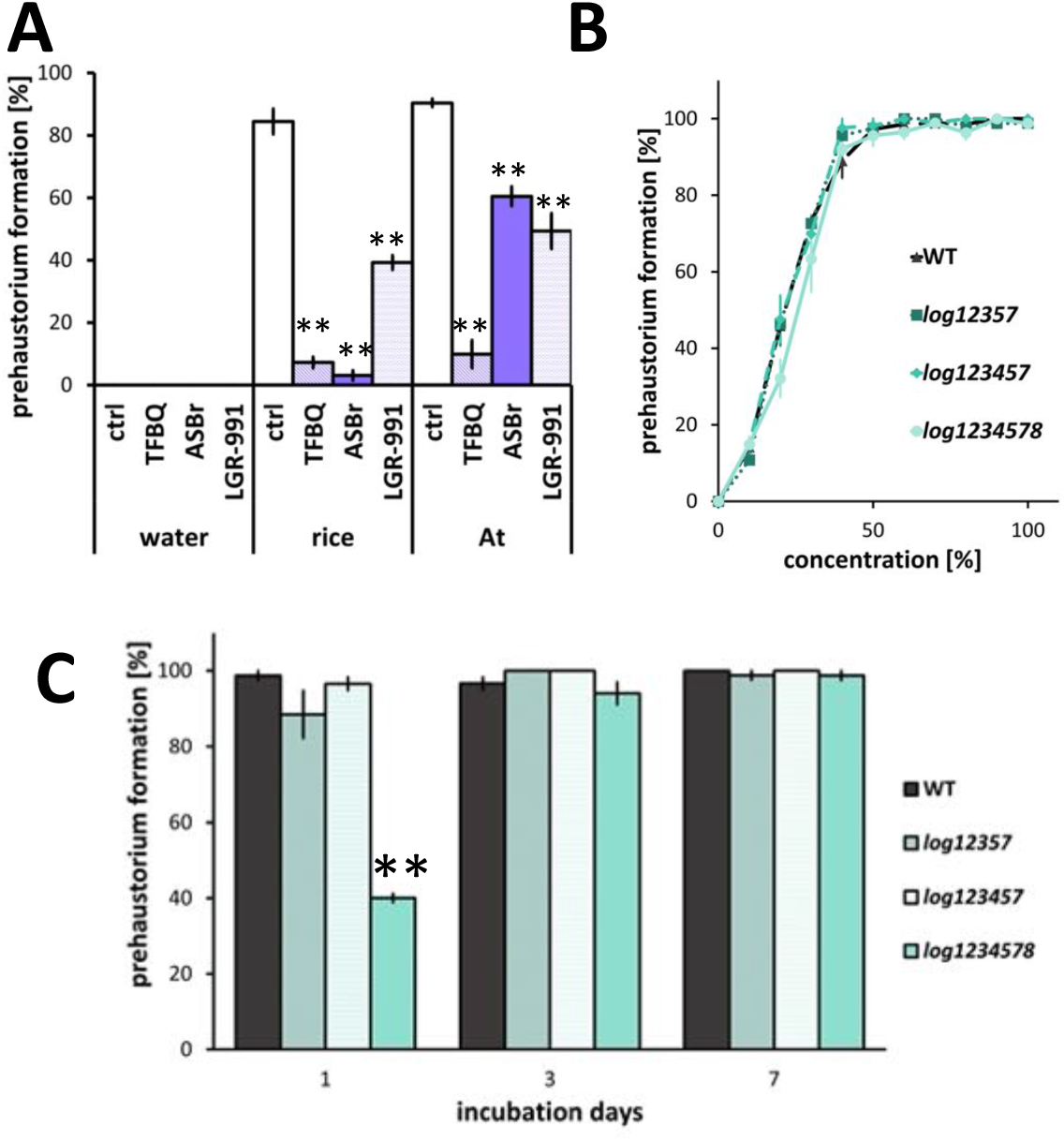
Prehaustorium-inducing activity of host plant exudates on *S. hermonthica*. A, The effect of inhibitors on *S. hermonthica* prehaustorium induction by rice or *A. thaliana* (At) exudates. TFBQ (20 µM), ASBr (10 µM), LGR-991 (10 µM). Plant exudates were diluted to a 40% concentration with water. B, C, Prehaustorium-inducing activities of root exudates from Arabidopsis *log12357, log123457* and *log1234578* mutants on *S. hermonthica*. B, Root exudates collected from *A. thaliana* after 7 days were diluted with water to the concentration indicated on the x-axis and tested for prehaustorium-inducing activity. C, Root exudates were collected from *A. thaliana* seedlings after incubation in water for 1 d, 3 d or 7 d and applied to *S. hermonthica* roots. The number of prehaustoria after a 24 h treatment was measured. Data represent the mean ± standard error (n=3). Asterisks indicate statistically significant differences compared to the control (*t*-test, **: *p*<0.01, *: 0.01<*p*<0.05).

To examine whether host root exudates contain cytokinins, we analyzed prehaustorium-inducing activities of *S. hermonthica* in the presence of root exudates from *A. thaliana* mutants defective in cytokinin biosynthesis (Fig. 5B). *A. thaliana* has nine *LOG* genes (*LOG1* to *LOG9*), of which LOG1 to LOG5, LOG7 and LOG8 have phosphoribohydrolase activity (Kuroha et al. 2009). Seedlings of the hextuple mutant *log123457* were reported to have reduced tZ and iP levels compared with WT seedlings (Tokunaga et al. 2012). In our study, we used *log12357, log123457* and *log1234578* mutants. Pentuple *log12357* and hextuple *log123457* did not show significant differences from WT in inducing prehaustorium formation in *S. hermonthica* (Figs. 5B and 5C). By contrast, the heptuple *log1234578* mutant had lower prehaustorium-inducing activity than the WT when the exudates were compared at a lower concentration (20-40% dilution) (Figs. 5B and 5C). These results suggest that cytokinin biosynthesis in the host contributes to HIF generation; however, since *log1234578* is reported to display severe growth defects (Tokunaga et al. 2012), the possibility that the reduction in prehaustorium-inducing activity was due to reduced root growth could not be eliminated.

### Gene expression induced by host root exudates

Finally, we analyzed the expression of the same genes analyzed in Fig. 4 but this time in response to host root exudates. We reasoned that if host root exudates contain a mixture of all types of HIFs, upregulation of any of these genes would also occur. Indeed, *ShQR2, ShPIRIN, ShYUC3, ShEXPB1* and two homologous chitinase genes were markedly upregulated at 1 and 6 h after rice exudate treatment (Fig. 6A). Although the response was not as extreme as in rice, treating *Arabidopsis* root exudates also upregulated this set of genes, particularly at 1h after treatment (Fig. 6A). Importantly, two cytokinin-responsive genes, *ShRR5* and *ShCKX2*, upregulated only by cytokinin *t*Z but not quinones or phenolics, were also upregulated by the rice and Arabidopsis root exudates after 1 and 6 h (Fig. 6B). This result indicates that host exudates trigger cytokinin responses in *S. hermonthica. ShIPT2*, a cytokinin biosynthesis gene, was upregulated by rice root exudate at 1-6 h. Similarly, *ShLOG5* was significantly upregulated at 1 h and to a greater extent at 6 h by rice root exudate. Both *ShIPT2* and *ShLOG5* showed slightly higher expression levels by *Arabidopsis* root exudates at 1 and 6 h, although without statistical significance. These results suggest that, unlike quinones or phenolic HIFs, host root exudates promote cytokinin biosynthesis and signaling (Fig. 6D). In summary, the expression of all prehaustorial marker genes and cytokinin-related genes was induced by host root exudates, suggesting the possibility that host root exudates contain a mixture of various HIFs.

**Fig. 6.**
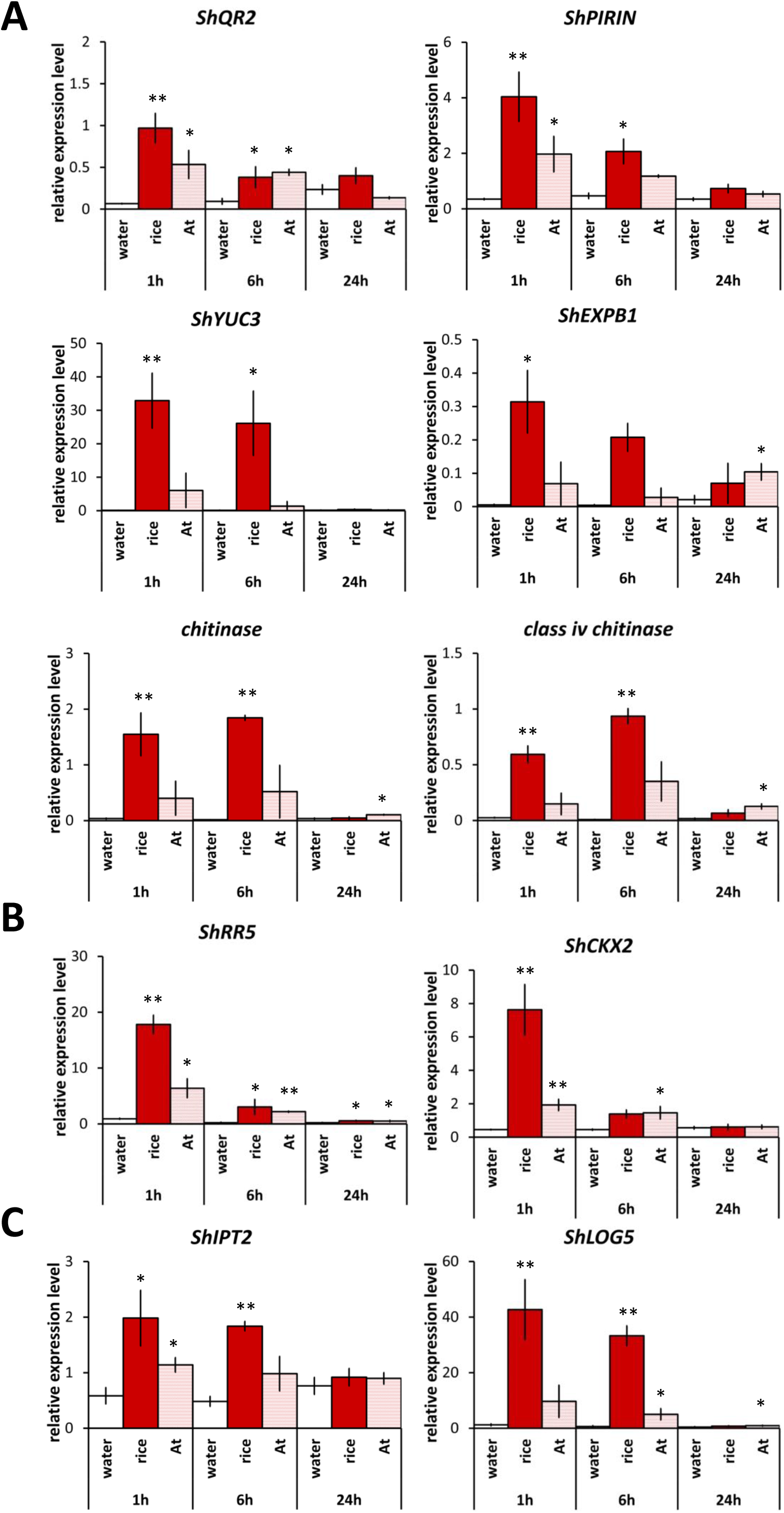
Expression analyses of genes involved in prehaustorium formation, cytokinin signaling and cytokinin biosynthesis in response to treatments with host root exudates. A, Expression of DMBQ marker genes (*ShQR2, ShPIRIN, ShYUC3, ShEXPB1*, chitinase and class iv chitinase). B, Expression of genes involved in cytokinin signaling (*ShRR5* and *ShCKX2*). C, Expression of genes involved in cytokinin biosynthesis (*ShLOG5* and *ShIPT2*). *S. hermonthica* was treated with rice or *A. thaliana* (At) exudates for 24 h. Data represent the mean ± standard error (n=3). Asterisks indicate significant differences compared to the control (*t*-test, **: *p*<0.01, *: 0.01<*p*<0.05).

## Discussion

### Cytokinins function in inducing terminal prehaustoria

Cytokinins were reported to induce prehaustorium formation in *P. ramosa* and *S. asiatica* (Goyet et al. 2017, Keyes et al. 2000). Our present work showed that cytokinins function as HIFs in *S. hermonthica* even at lower concentration than the well-known HIF, DMBQ (Fig. 7A). Although they are differed in photosynthetic ability, *e*.*g. Striga* spp. are hemiparasites and *P. ramosa* is a holoparasite, these plants are all obligate parasitic plants that primarily form terminal haustoria at the radicle tip resulting from deformation of root apical meristems. We also found in this study that *P. japonicum* does not form prehaustoria in response to cytokinin. *P. japonicum* is a facultative hemiparasitic plant that forms lateral haustoria on the sides of roots without disturbing root growth. Previously, *T. versicolor*, another facultative parasite, was reported to form prehaustoria-like structures with cytokinin (BA) treatment (Wrobel and Yoder 2001). However, BA treatment of *T. versicolor* roots caused root tip swelling and hair proliferation, which resembles the morphology of a terminal prehaustorium. When the roots were transferred to media without BA after 4 h of BA treatment, the morphology of prehaustoria resembled the lateral prehaustoria formed in DMBQ-treated *T. versicolor* (Wrobel and Yoder 2001). Thus, we infer that cytokinin is an inducer of terminal prehaustoria but not lateral prehaustoria. Accumulative knowledge from previous studies suggested that quinones, phenolic compounds and flavonoids are common HIFs for hemiparasitic Orobanchaceae, including *Striga, Phtheirospermum, Triphysaria, Castilleja* and *Agalinis* (Goyet et al. 2019). On the other hand, holoparasitic *Phelipanche* and *Orobanche* are not highly responsive to those compounds (Goyet et al. 2017, Fernandez-Aparicio et al. 2021). Therefore, it was hypothesized that species specificity of HIFs could be determined by the photosynthesis ability. Our study adds a new insight for specificities of cytokinin-type HIFs as those could be determined by haustorial morphology, that is either terminal or lateral. Because only a few parasite species were analyzed for various HIFs so far, it would be necessary to analyze more parasite species to validate this hypothesis. Interaction between cytokinin and auxin may explain different HIF activities to *S. hermonthica* and *P. japonicum*. During early prehaustorium formation in *P. japonicum*, auxin biosynthesis mediated by YUC3 at the prehaustorium apex is crucial for prehaustorium development (Ishida et al. 2016). External application of auxin also increases the number of prehaustoria induced by DMBQ in *P. japonicum* (Fig. S7). Cytokinin was shown to suppress asymmetric cell division promoted by auxin and inhibits lateral root initiation (Laplaze et al. 2007). Therefore, exogenous cytokinin may repress auxin signaling and inhibit prehaustorium formation in *P. japonicum*. In addition, cytokinins were reported to be negative regulators of primary root growth by modulating root meristems (Werner et al. 2010). In this regard, cytokinins are proposed to suppress root growth by terminating root meristematic activity, a role important for forming terminal prehaustoria that require the deformation of a growing root tip.

**Fig. 7.**
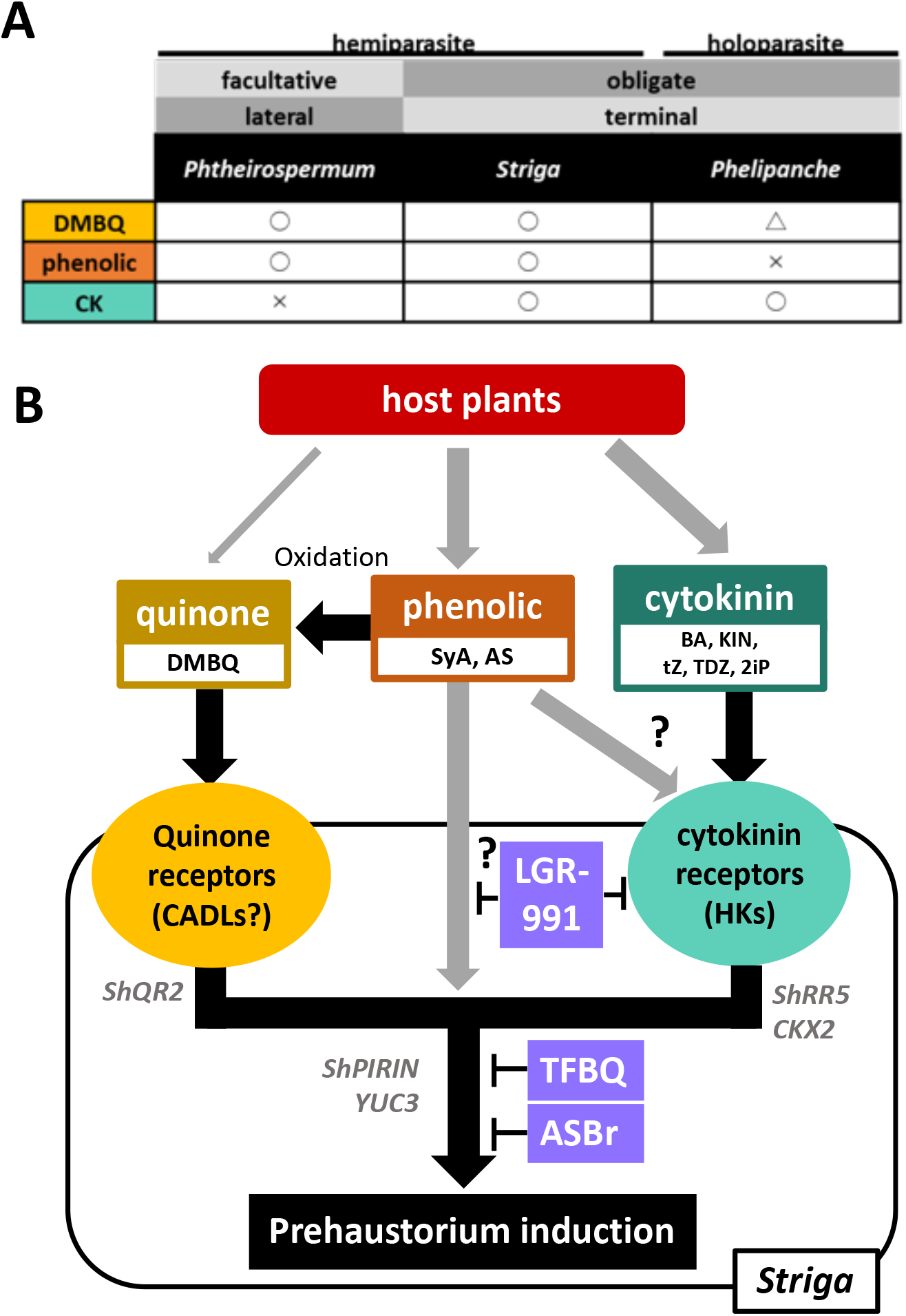
Classification and model of the HIF responses in Orobanchaceae. A, The classification of Orobanchaceae parasitic plants and the prehaustorium inducing activities of DMBQ, phenolics or cytokinin. B, A model for the pathway of prehaustorium induction in *Striga*.

### The relationships of signaling pathways for prehaustorium induction by HIFs

The finding that *S. hermonthica* prehaustoria can be induced by three types of HIFs enabled us to investigate the relationships among the signaling pathways induced by each HIF. The conversion from phenolic HIFs to quinones by oxidation was confirmed by *in vitro* experiments (Kim et al., 1998). Thus, we hypothesized that the signaling pathway for phenolics AS or SyA overlaps with that of DMBQ. Indeed, several ROS inhibitors similarly inhibit prehaustorium formation in *S. hermonthica* induced by SyA and DMBQ (Wada et al. 2019). This study showed that TFBQ and ASBr suppressed prehaustorium formation induced by AS, SyA or DMBQ (Figs. 3D and 3E). TFBQ is a known DMBQ inhibitor of prehaustorium induction in parasitic plants (Smith et al. 1996, Wang et al. 2019) and inhibits DMBQ-induced Ca^2+^ uptake mediated by CARD1 receptor kinase in Arabidopsis (Laohavisit et al. 2020). Although CARD1 receptor kinase does not respond to SyA in Arabidopsis, parasitic plants may convert SyA or AS to DMBQ as a direct signaling molecule and, therefore, prehaustoria induced by SyA or AS were effectively inhibited by TFBQ. ASBr was reported to be an inhibitor of VirA, a sensor of plant-derived AS in *Agrobacterium* (Hess et al. 1991). To our knowledge, this was the first report of ASBr also inhibiting AS-derived signaling in plants. Interestingly, ASBr also suppressed DMBQ-induced prehaustoria in *S. hermonthica*, although ASBr was believed to be a competitive inhibitor for AS. TFBQ and ASBr possess electronegative halogen group(s) that tend to gain an electron and, thus, show high reactivity. Because the redox status is important for prehaustorium induction (Smith et al. 1996, Wada et al. 2019), these inhibitors may inhibit prehaustorium formation by disturbing the redox status. Indeed, TFBQ and ASBr inhibited prehaustorium formation by cytokinins, indicating that these inhibitors target the common downstream signaling pathways of quinone, phenolic and cytokinin-types of HIFs.

LGR-991, a competitive inhibitor of cytokinin receptors, effectively inhibited cytokinin-induced prehaustorium formation and moderately inhibited AS- or SyA-induced but not DMBQ-induced prehaustoria in *S. hermonthica*. These results suggest that the phenolic signaling and cytokinin signaling pathways overlap partially, but the DMBQ pathway does not. Because LGR-991 directly binds cytokinin receptors (Nisler et al. 2010), phenolic signals may be located upstream of the cytokinin signaling pathway (Fig. 7B). However, phenolic HIF treatments did not increase cytokinin-responsive genes in *S. hermonthica* (Fig. 4B). Possibly, cytokinin activation after SyA or AS treatments is tentative or local and, therefore, not detected in RT-qPCR experiments. Alternatively, LGR-991 inhibits signaling molecules involved in phenolic HIF signaling apart from cytokinin receptors. Because the *ShQR2* and *ShEXPB1* genes were upregulated by DMBQ and SyA but not by cytokinin, these genes are specific to phenolic/quinone pathway and activated upstream of quinone-cytokinin converging route (Fig. 7). On the other hand, *PIRIN, YUC3* and chitinases are upregulated by all three types of HIFs. This reflects a converged transcript activation pathway in response to different HIFs in inducing prehaustorium. It is noteworthy that, unlike *S. hermonthica*, the DMBQ-induced prehaustoria in *P. japonicum* can also be inhibited by LGR-991 treatment. Thus, the DMBQ-dependent signal in *P. japonicum* may differ from that of *S. hermonthica*.

### HIFs in host plant root exudates

DMBQ was first isolated from *Sorghum* extract (Chang and Lynn 1987) and is considered a major host-derived HIF. However, the amount of DMBQ in Arabidopsis root exudates is too low to account for its HIF activity (Wang et al. 2020). In this study, prehaustorium induction by rice and Arabidopsis root exudates was reduced by TFBQ, ASBr and LGR-991 treatments (Fig. 5A). Because LGR-991 inhibited prehaustorium induction, the primary HIFs in host root exudates may be phenolic or cytokinin-type HIFs. Furthermore, when we used exudates from the Arabidopsis *log* mutants, there was only a subtle change in prehaustorium induction activity, suggesting that Arabidopsis root exudates contain phenolic-type HIFs (Figs. 5B and 5C). Also, DMBQ or SyA upregulated genes that are highly induced by host exudates. However, *S. hermonthica* treated with host root exudates also expressed cytokinin biosynthesis and responsive genes (Figs. 6B and 6C). This result is consistent with previous findings that *P. ramosa* expresses cytokinin-responsive genes during prehaustorium formation induced by host exudates (Goyet et al. 2017). Endogenous cytokinin production in parasitic plants upon host root exudate treatment may have function in further promoting prehaustorium formation. *Arabidopsis* root exudates have not been reported to contain cytokinins, but several reports have shown that cytokinins are present in rice exudates (Murofushi et al. 1983, Soejima et al. 1992, 1995). Plant root exudates are known to contain various phenolic acids, including ferulic acid and sinapic acid that have haustorium inducing activity (Narasimhan et al. 2003). Taken together, these observations suggest that the host root exudates, at least in rice, are likely to contain phenolic and cytokinin-type types of HIFs.

In conclusion, we have shown that nano molar ranges of cytokinins can induce prehaustoria in *Striga* but not *P. japonicum*, which may influence host sensitivity and selectively of the weedy parasitic plants. Presence of multiple HIFs pathways may imply complexity and robustness of prehaustorium induction signaling. This can be taken into account to develop *Striga* control methods. Determining exact HIFs involved in host exudates and each signaling component in parasitic plants remains as future research objectives. Combination of genetic and biochemical analysis would dissect host-parasite signaling interaction.

## Materials and Methods

### Plant materials

*S. hermonthica* seeds were sterilized with 20% (v/v) commercial bleach solution (Kao) 5 times and then washed with sterile water 10 times. The sterilized seeds were transferred to 9 cm petri dishes filled with moistened Whatman Glass Microfiber Filters GF/A (GE Healthcare, Little Chalfont) and incubated at 25 °C for 4-7 days in dark conditions. To germinate *S. hermonthica*, seeds were treated with 10 nM strigol (Hirayama and Mori 1999) at 25 °C for 24 h in dark conditions.

*P. japonicum* wild type seeds (ecotype: Okayama) were sterilized with 10% commercial bleach solution 5 times and washed with sterile water 5 times. Sterilized seeds were sown on 1/2 MS (Murashige and Skoog Plant Salt Mixture, Wako Pure Chemical Co.) media (pH 5.8) with 0.8% (w/v) agar and 1% (w/v) sucrose and kept at 4 °C in the dark for 3 days. The plates were moved to a controlled temperature chamber set at 25 °C with a photoperiod of 16 h light / 8 h dark and a light intensity of 90 µmol m^−2^ s^−1^.

*A. thaliana* ecotype Columbia (Col-0) was used as the wild type. The *log12357, log123457* and *log1234578* mutants were previously reported (Kuroha et al. 2009, Tokunaga et al. 2012). Their seeds were sterilized with 5% (v/v) commercial bleach solution and washed with sterile water 5 times. Sterilized seeds were sown on 1/2 MS media with 0.8% (w/v) agar and 1% (w/v) sucrose, kept at 4 °C in the dark for 3 days, and grown at 22 °C with a photoperiod of 16 h light / 8 h dark and a light intensity of 69 µmol m^−2^ s^−1^.

Rice (*Oryza sativa* cv. Koshihikari) seeds were washed with 70% (v/v) ethanol for 5 minutes and then sterilized with a 20% (v/v) commercial bleach solution for 30 minutes with stirring. After sterilization, seeds were rinsed at least 5 times with sterilized water. For exudate sampling, rice seeds were transferred to wet filter paper (Advantec Co.) in 9 cm petri dishes and incubated at 25 °C for 7-10 days under 16 h light / 8 h dark, 90 µmol m^−2^ s^−1^ conditions.

### Chemicals

DMBQ (Sigma-Aldrich), acetosyringone (Tokyo Chemical Industry Co.), syringic acid (Sigma-Aldrich), kinetin (Sigma-Aldrich), 2iP, TFBQ (Sigma-Aldrich), ASBr (Nacalai Tesque), PI-55 (Spíchal et al. 2009), LGR-991 (Nisler et al. 2010) were dissolved in dimethyl sulfoxide (DMSO, Nacalai Tesque) as 10 mM stocks. BA (Wako Pure Chemical Co.), *trans*-zeatin (Wako Pure Chemical Co.) and thidiazuron (Wako Pure Chemical Co.) were dissolved in sterile water as 1 mg/mL stocks. These stocks were diluted with sterile water in each experiment.

### Collection of plant root exudates

Five *A. thaliana* seedlings grown in 1/2 MS agar plates for 10-14 days were transferred to 6-well plates (Iwaki) containing 5 ml sterile water/well and incubated at 25 °C for 1, 3 or 7 days in the dark. Single rice seedlings of 7-10 days old were transferred into each of the wells and incubated at 25 °C for 7 days in the dark. The water in the well was collected and used as the source of plant exudates.

### Plant growth experiments and prehaustorium induction assay

Germinated *S. hermonthica* seedlings were transferred to 96-well plates (Iwaki) containing 100 µL water per well with or without chemicals and incubated at 25 °C in the dark. Approximately 20-40 seedlings were placed in each well. After 24 h, the number of prehaustoria was counted using a stereo microscope. Prehaustorium formation was calculated as a percentage of the number of *S. hermonthica* seedlings forming prehaustoria divided by the number of total *S. hermonthica* seedlings in each well.

*P. japonicum*, grown in 1/2 MS agar plates with 1% (w/v) sucrose for 7 days, were transferred to 1% (w/v) agar plates containing DMBQ or cytokinins. Plates were incubated at 25 °C with a photoperiod of 16 h light / 8 h dark, 90 µmol m^−2^ s^−1^. Lateral root number, root length and prehaustorium number were measured after 7 days for the data presented Figs. S1A-C. For Figs. S4B-D, seedlings of *P. japonicum*, grown in 1/2 MS agar plates with 1% (w/v) sucrose for 10 days, were transferred to 1% (w/v) agar plates containing the indicated chemicals and incubated for 7 days with a photoperiod of 16 h / per 8 h dark, 90 µmol m^−2^ s^−1^ before measuring lateral root number, root length and hypocotyl length.

For the prehaustorium induction assay in *P. japonicum* (Fig. S5A), seedlings, grown in 1/2 MS agar plates with 1% (w/v) sucrose for 7 days, were transferred to 6-well plates. Ten seedlings and 5 mL water were placed in each well and incubated at 25 °C overnight in the dark. *P. japonicum* were then moved to another 6-well plate containing 5 mL water and selected HIFs or inhibitors in each well. Plates were incubated at 25 °C for 5 days in the dark before measuring prehaustorium formation.

### RT-qPCR

RNA was extracted from *S. hermonthica* treated with water or HIFs for 1, 6 or 24 h using an RNeasy Plant Mini Kit (Qiagen) according to the manufacturer’s instructions. Genomic DNA was removed from samples by RNase-free DNase I (Qiagen) treatment. The total RNA concentration was measured using a NanoDrop ND-1000 (NanoDrop Technologies). Extracted RNAs (500 ng) were reverse transcribed to cDNA using the ReverTra Ace qPCR RT Master Mix (Toyobo) according to the manufacturer’s instructions. cDNA was diluted 10 times, and 2 µL was used as the template in 20 µl of THUNDERBIRD SYBR qPCR Mix (Toyobo) with 0.3 mM of each primer. Quantification of RT-qPCR products was carried out on a CFX-Connect (Bio-Rad) according to the manufacturer’s instructions using the standard curve method. Actin was used as an internal control. Target genes in *S. hermonthica* were searched by blastp using the corresponding *A. thaliana* genes and the assembled *S. hermonthica* transcriptome (Yoshida et al. 2019) as the queries. The primer sets used in this study is listed in Table S1.

## Data Availability

The data underlying this article are available in the article and in the online supplementary material.

## Funding

This work was partly supported by KAKENHI grant numbers JP19K16169 to SC, JP19K22432, JP21H02506 and JP20H05909, and JST PRESTO grant number JPMJPR194D to SY. This work was partially supported by the Asahi Glass Foundation.

## Disclosures

Conflicts of interest: No conflicts of interest are declared.

## Acknowledgements

The authors thank Prof. A. G. Babiker (Environment and Natural Resources and Desertification Research Institute, Sudan) and Emer. Prof. Kenji Mori (The University of Tokyo) for kindly providing *Striga hermonthica* seeds and strigol, respectively. The inhibitors PI-55 and LGR-991 were provided by Dr. Lukáš Spíchal (Palacký University, Czech Republic) and the *log12357, log123457* and *log1234578* mutants were a gift from Drs. Takatoshi Kiba and Hitoshi Sakakibara (Nagoya University). NA was supported by the NAIST University Fellowships for the Creation of Innovation in Science and Technology from the Nara Institute of Science and Technology.

## Author Contributions (optional)

NA and SY conceived this study. NA and SC performed the experiments. NA, SC and SY wrote the manuscript.

## Legends to Figures

**Fig. S1.**
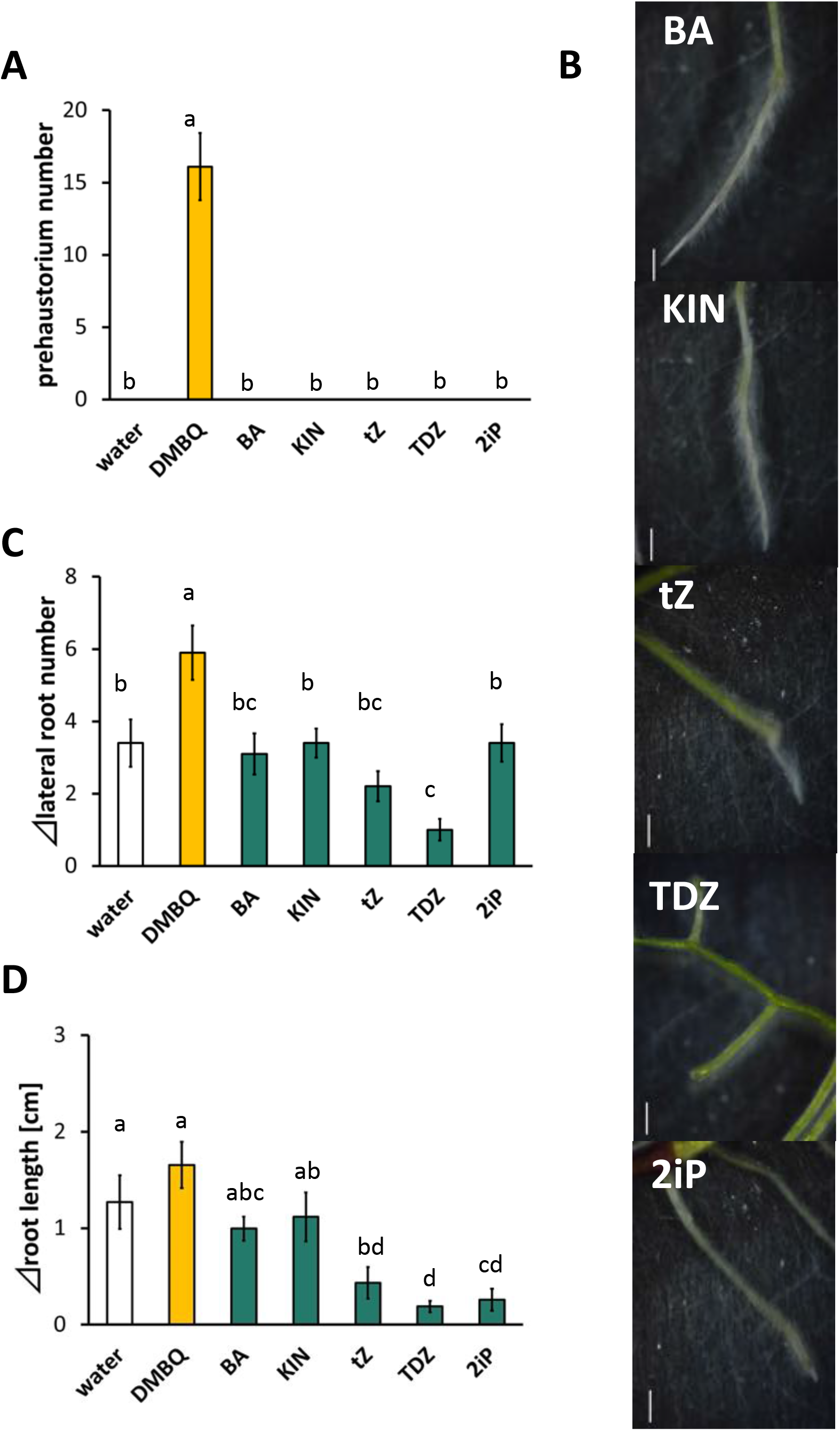
Prehaustorium induction and root growth in *P. japonicum* after cytokinin treatment. A, The effects of cytokinins on *P. japonicum* prehaustorium formation. B, Changes in *P. japonicum* root morphology after treatment with cytokinins. C, The effect of cytokinins on *P. japonicum* lateral root initiation. D, The effect of cytokinins on *P. japonicum* root elongation. *P. japonicum* seedlings were incubated for 7 days on agar plates containing 500 nM BA, KIN, tZ, TDZ or 2iP. Scale bars=1 mm. Data represent the mean ± standard error (n=10). Letters indicate significant differences (Tukey’s test, *p*<0.05).

**Fig. S2.**
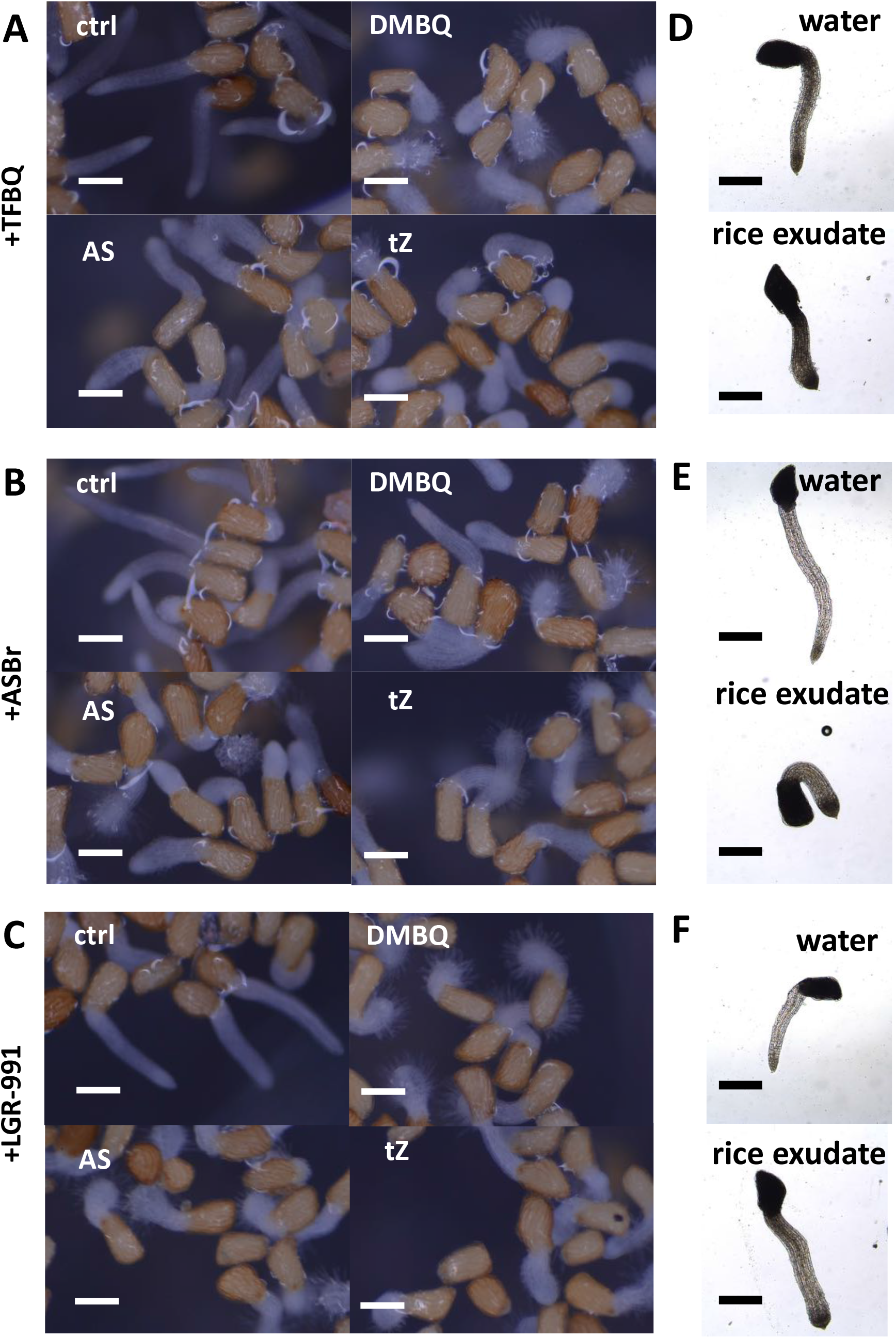
The effects of chemical inhibitors of HIFs on *S. hermonthica* radicals during prehaustorium formation. A-C, *S. hermonthica* seedlings after treatment with DMBQ, AS or tZ in the presence of TFBQ (A), ASBr (B) or LGR-991 (C). Inhibitors only (ctrl) were used as control to reflect their effects on root growth. D-F, Effects of TFBQ (D), ASBr (E) and LGR-991 (F) on rice exudates in inducing prehaustorium formation. 20 µM TFBQ 10 µM ASBr (10 µM) and 10 µM LGR-991 were used. Scale bars: 100 µm for A, B, C, 500 μm for D, E, F.

**Fig. S3.**
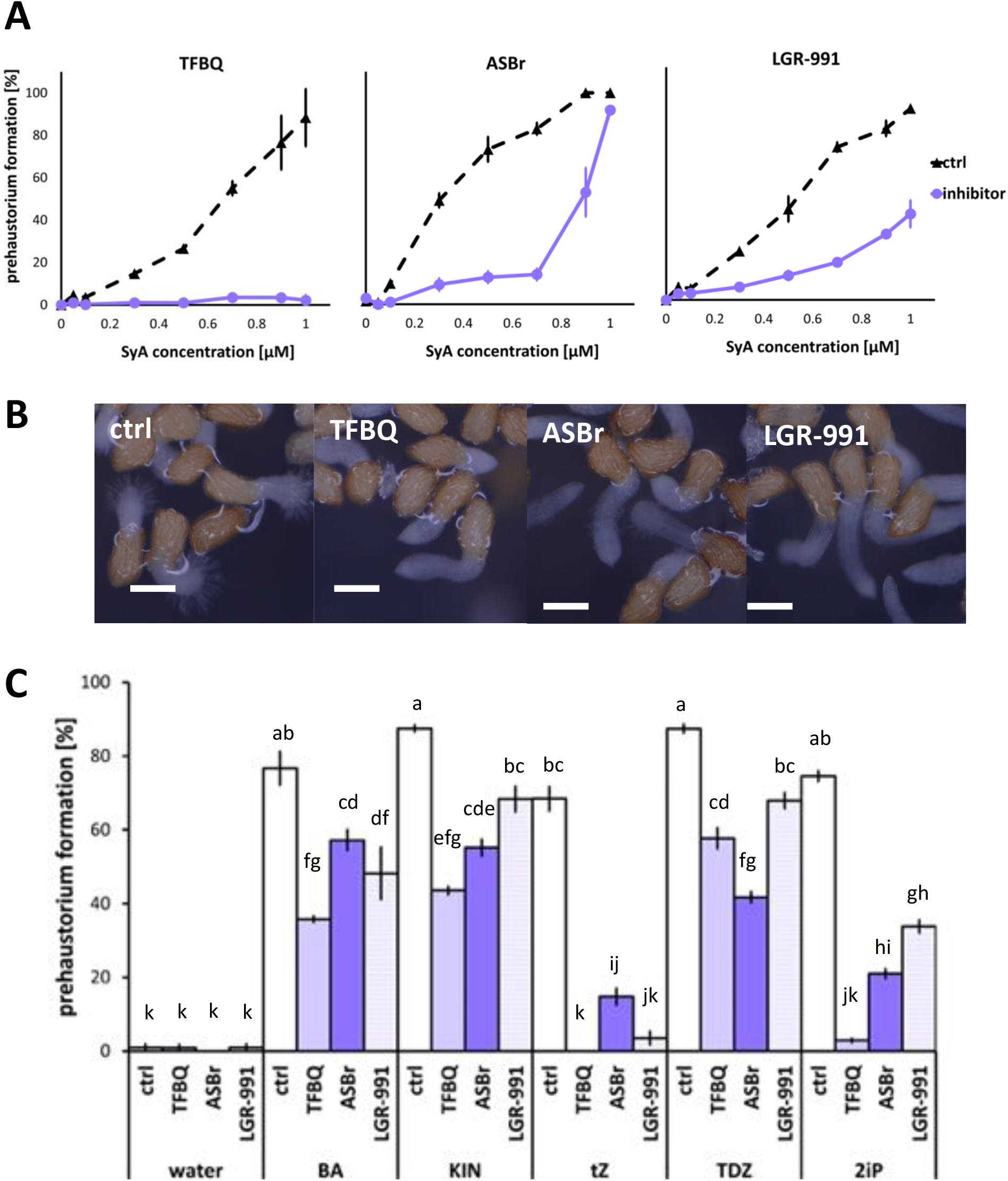
The effects of chemical inhibitors of HIFs in the presence of SyA or cytokinins on prehaustorium formation in *S. hermonthica*. A, The effects of TFBQ (20 µM), ASBr (10 µM) and LGR-991 (10 µM) on *S. hermonthica* prehaustorium induction with SyA treatment. B, Images of *S. hermonthica* seedlings treated with SyA in the presence of TFBQ, ASBr or LGR-991. SyA only (ctrl) was used as the control. C, The effects of TFBQ (100 µM), ASBr (10 µM) and LGR-991 (1 µM) on *S. hermonthica* prehaustorium formation induced by BA (300 nM), KIN (500 nM), tZ (25 nM), TDZ (25 nM) or 2iP (100 nM). Data represent the mean ± standard error (n=3). Letters indicate significant differences (Tukey’s test, *p*<0.05). Scale bars=100 µm.

**Fig. S4.**
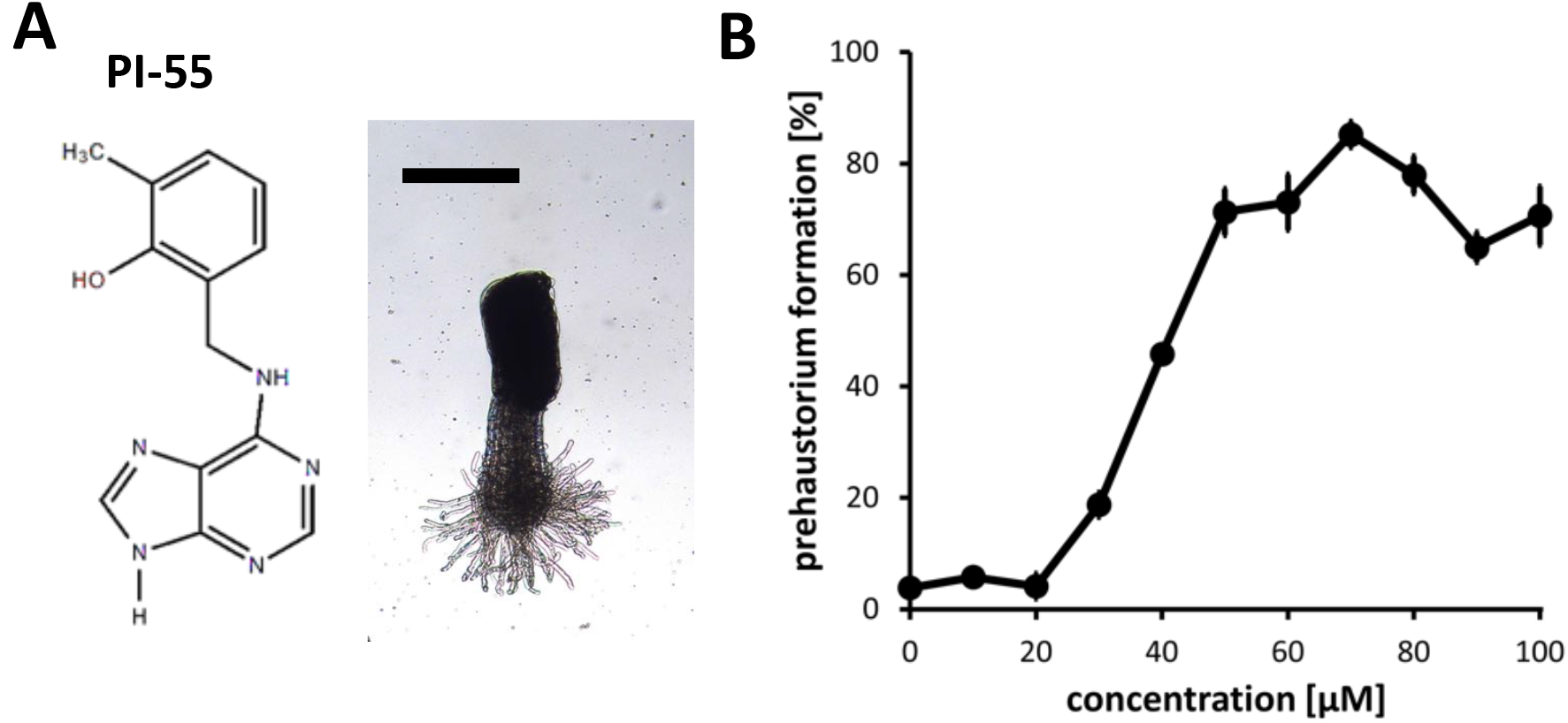
Cytokinin inhibitors PI-55 and LGR-991 can induce prehaustoria in *S. hermonthica*. A, The chemical structure of PI-55 (left) and morphology of prehaustoria induced by 10 µM PI-55 in *S. hermonthica* (right). Scale bar = 1 mm. B, LGR-991 induced prehaustoria in *S. hermonthica* at concentrations higher than 20 µM. *S. hermonthica* seedlings were exposed to HIFs and cytokinin inhibitors for 24 h. Data represent the mean ± standard error (n=3).

**Fig. S5.**
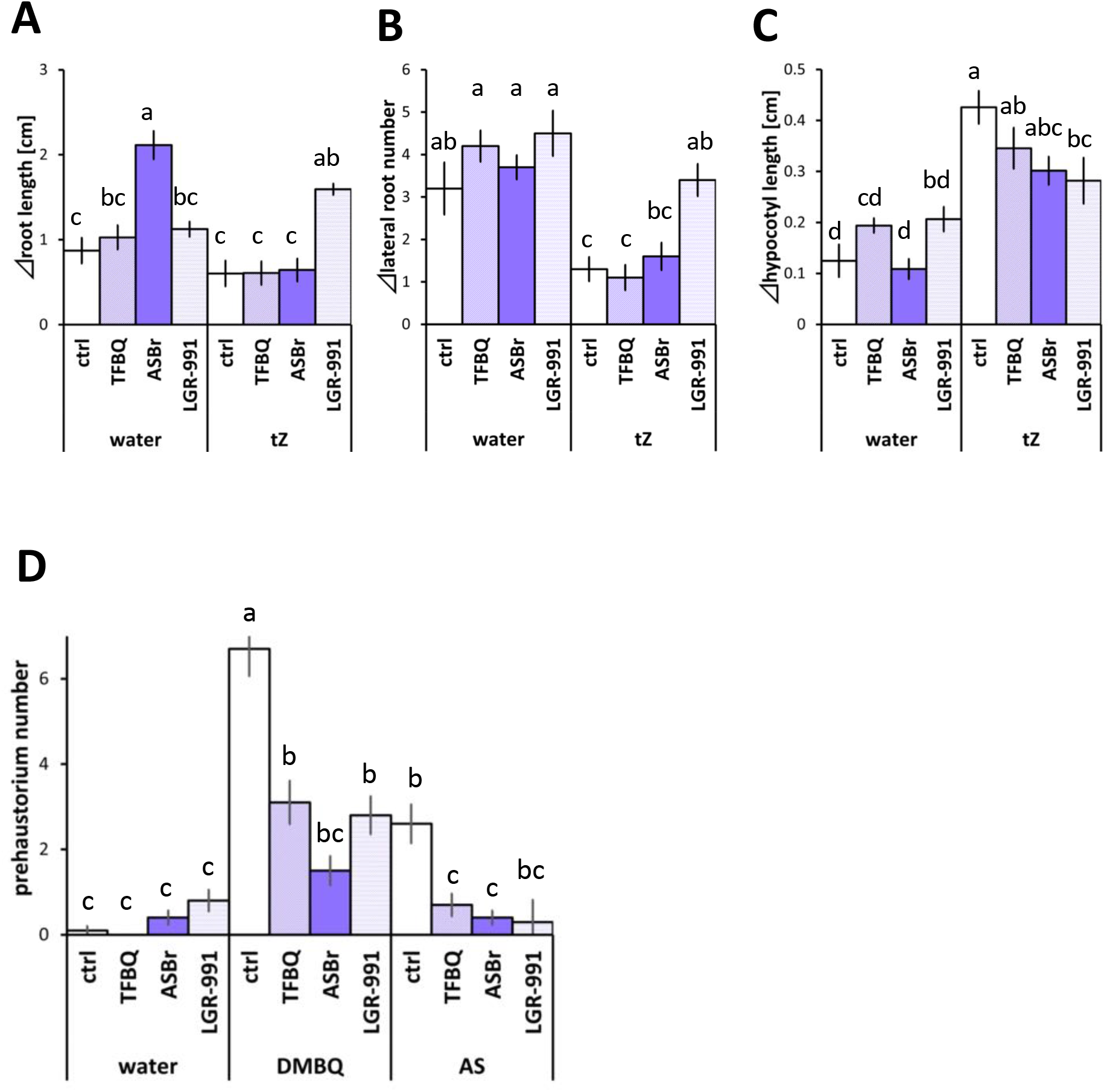
The effect of inhibitors on prehaustorium induction and cytokinin signaling in *P. japonicum*. A, The effect of TFBQ (20 µM), ASBr and LGR-991 (10 µM) on *P. japonicum* prehaustorium induction by DMBQ (10 µM) or AS (10 µM). *P. japonicum* seedlings were incubated for 5 days in HIFs and inhibitor solutions. B-C, Effects of TFBQ (20 µM), ASBr and LGR-991 (5 µM) on cytokinin signaling induced by tZ (500 nM) in *P. japonicum*. B, Effects of inhibitors on root elongation suppressed by tZ. C, Effects of inhibitors on lateral root initiation suppressed by tZ. D, Effects of inhibitors on hypocotyl growth promoted by tZ. Data represent the mean ± standard error (n=10). Letters indicate significant differences (Tukey’s test, *p*<0.05).

**Fig. S6.**
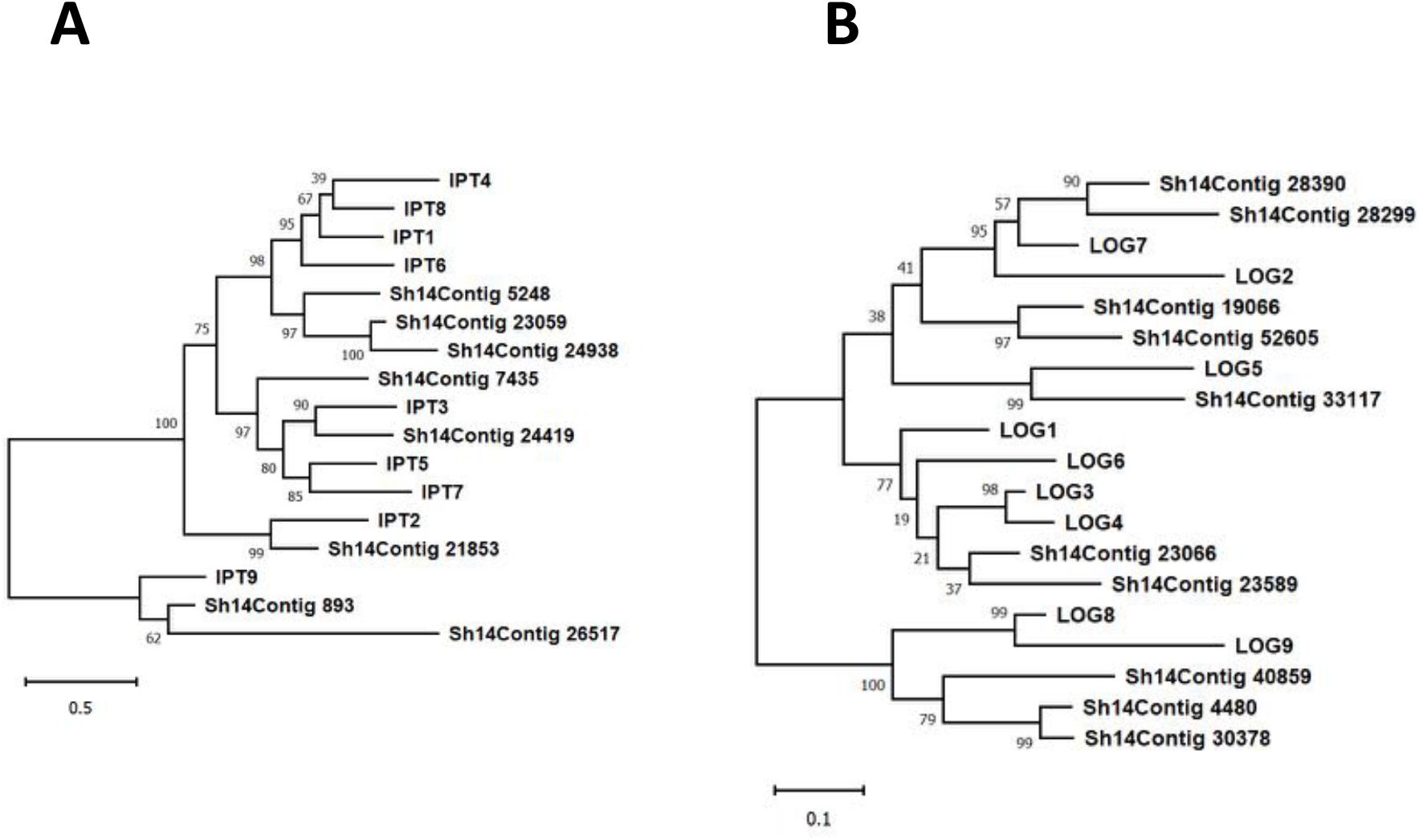
Phylogenetic trees of cytokinin biosynthesis genes. A, Phylogenetic tree of IPT genes in *A. thaliana* and *S. hermonthica*. B, Phylogenetic tree of LOG genes in *A. thaliana* and *S. hermonthica*.

**Fig. S7.**
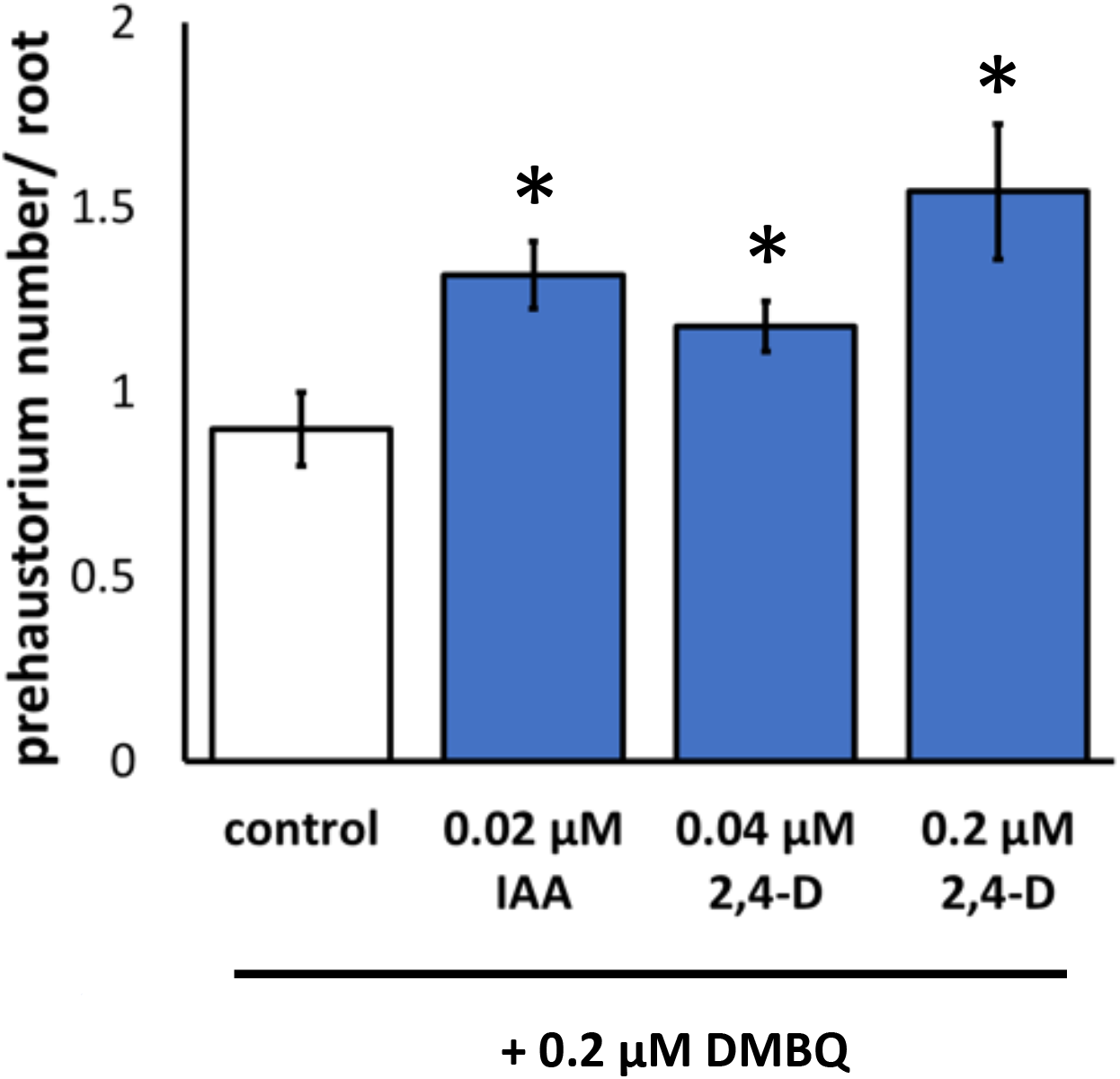
The effect of auxin on prehaustorium formation in *P. japonicum*. A, The effect of auxin on prehaustorium induction by DMBQ. *P. japonicum* seedlings were incubated for 4 days on agar plates containing DMBQ (0.2 µM) and Indole-3-acetic acid (IAA) or 2,4-Dichlorophenoxyacetic acid (2,4-D). Data represent the mean ± standard error (n=22). Asterisks indicate statistically significant differences compared to the control (t-test, p<0.05).

**Table S1.**
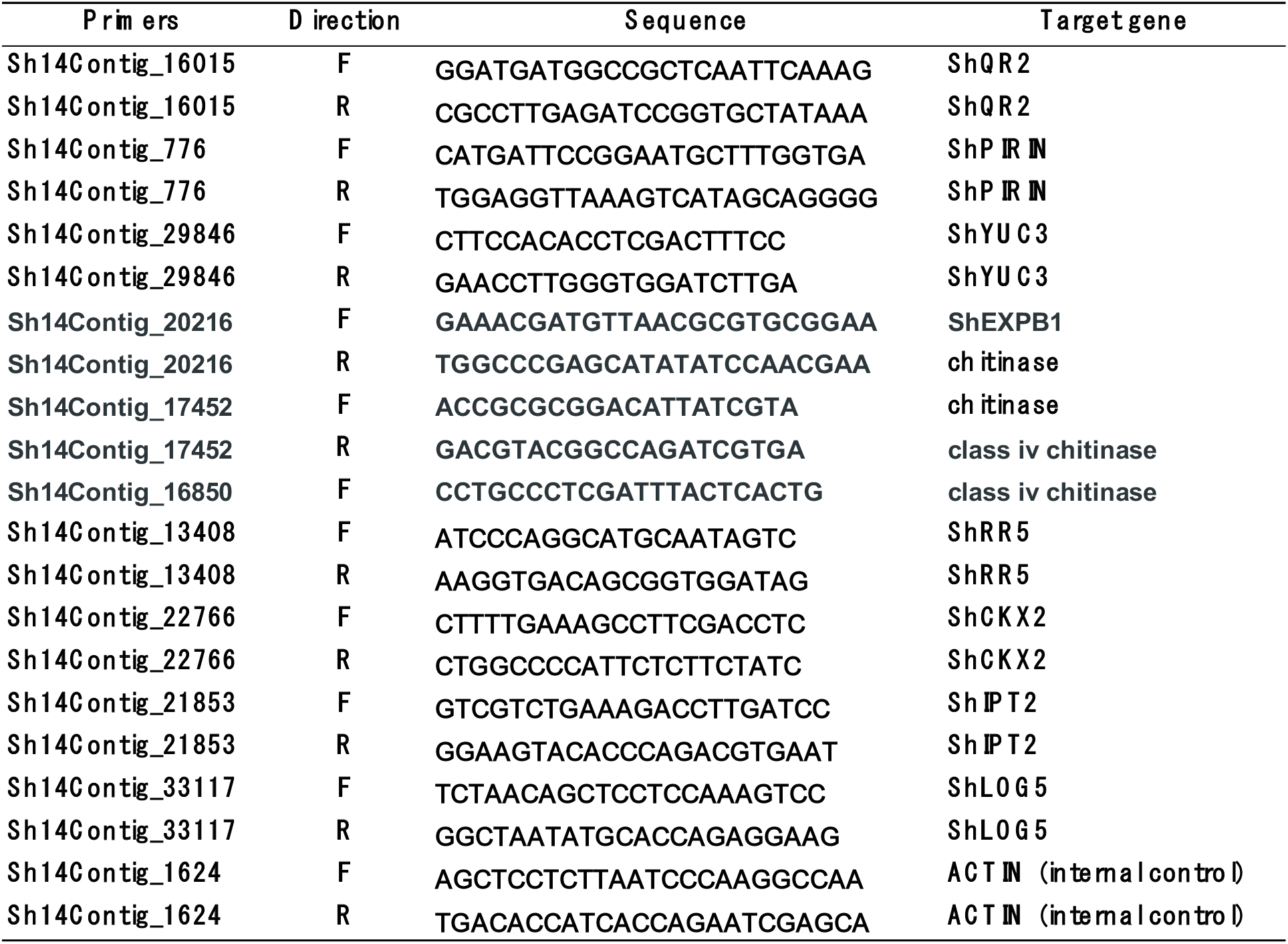
Primer list used in this study

## Notes

### Competing Interest Statement

The authors have declared no competing interest.

