## Supplemental figures for "Cytokinins induce prehaustoria coordinately with quinone and phenolic signals in the parasitic plant *Striga hermonthica*"

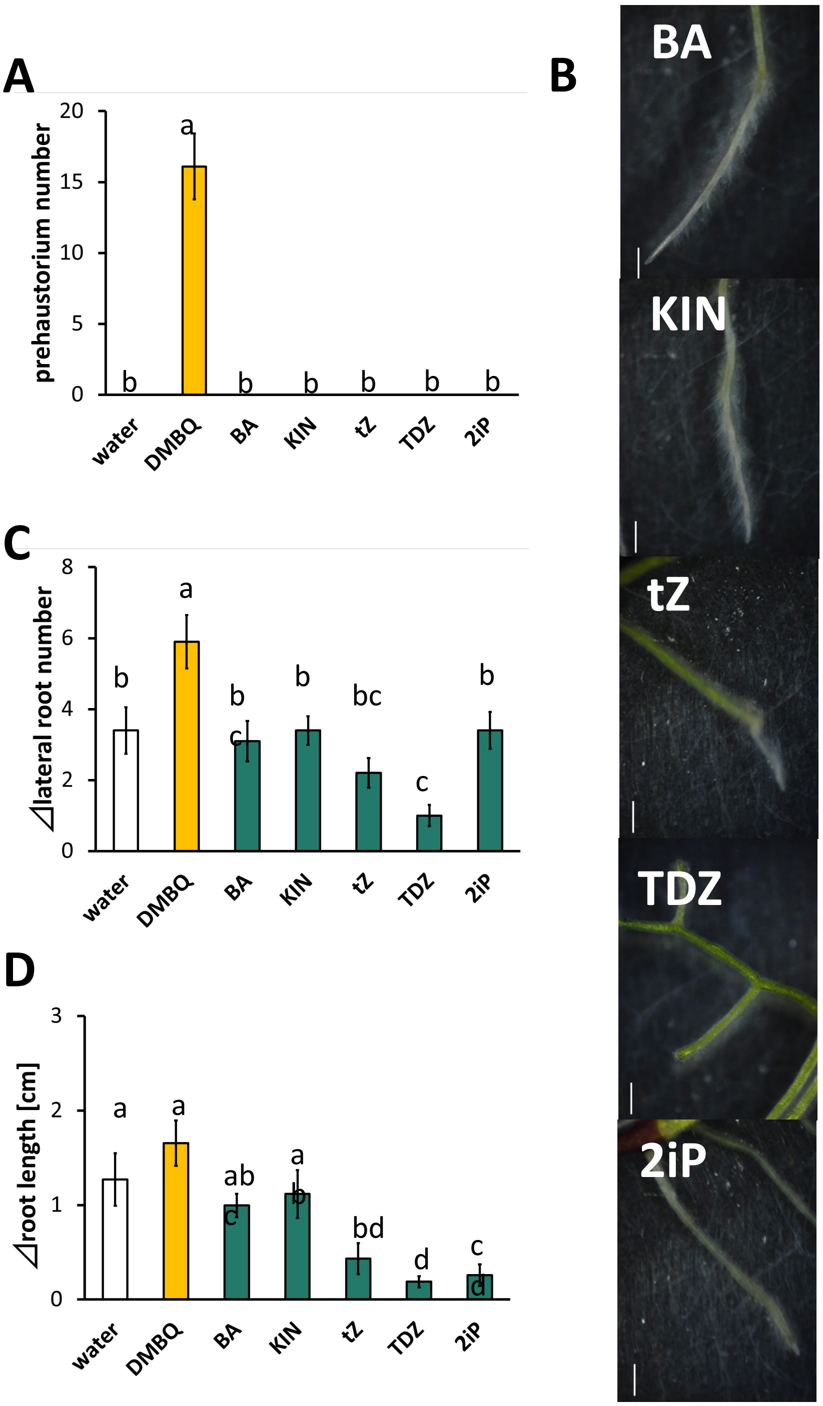


**Fig. S1** Prehaustorium induction and root growth in *P. japonicum* after cytokinin treatment.

A, The effects of cytokinins on *P. japonicum* prehaustorium formation. B, Changes in *P. japonicum* root morphology after treatment with cytokinins. C, The effect of cytokinins on *P. japonicum* lateral root initiation. D, The effect of cytokinins on *P. japonicum* root elongation. *P. japonicum* seedlings were incubated for 7 days on agar plates containing 500 nM BA, KIN, tZ, TDZ or 2iP. Scale bars=1 mm. Data represent the mean ± standard error (n=10). Letters indicate significant differences (Tukey's test, $p$<0.05).

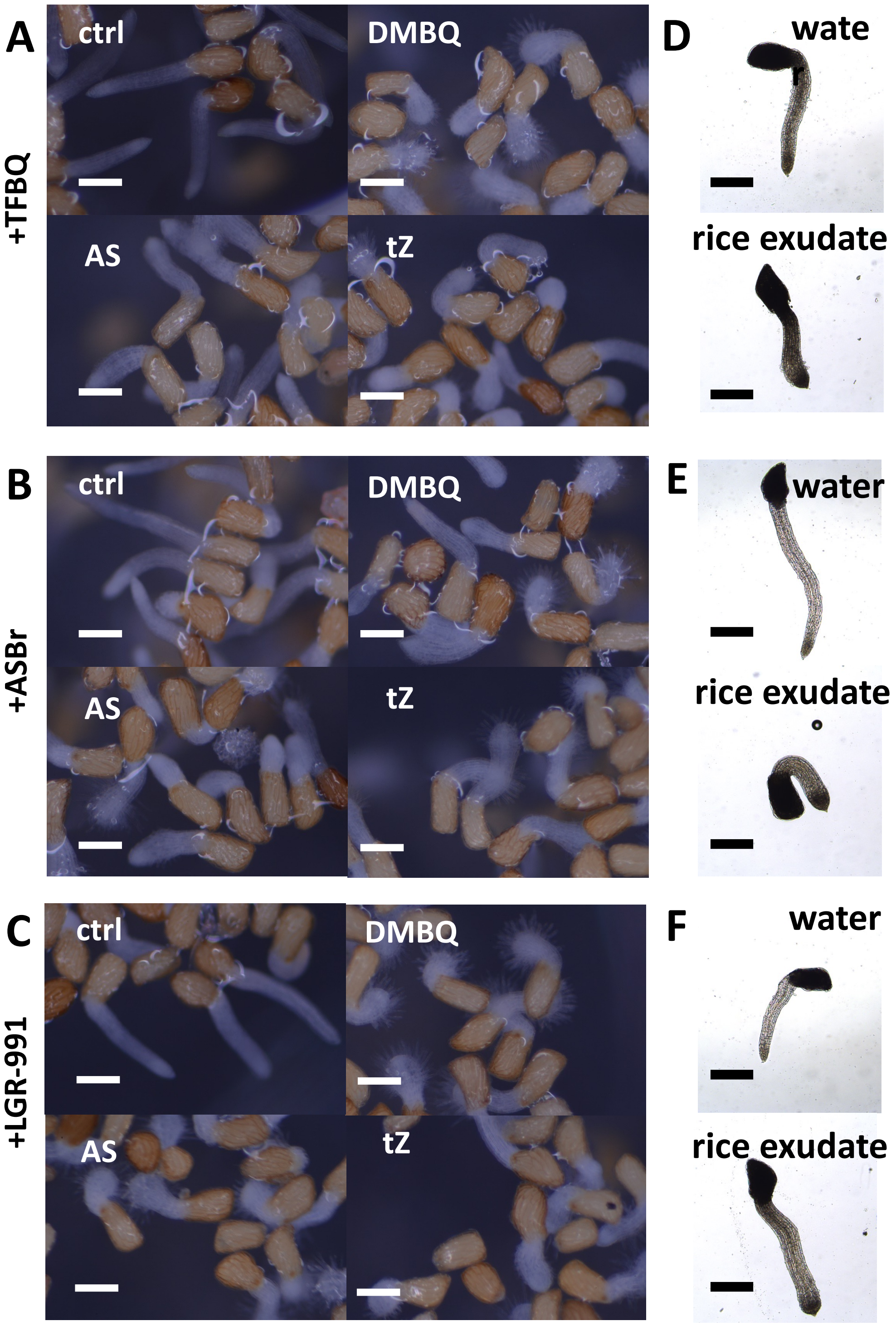


**Figure S2** The effects of chemical inhibitors of HIFs on *S. hermonthica* radicals during prehaustorium formation.

A-C, *S. hermonthica* seedlings after treatment with DMBQ, AS or tZ in the presence of TFBQ (A), ASBr (B) or LGR-991 (C). Inhibitors only (ctrl) were used as control to reflect their effects on root growth. D-F, Effects of TFBQ (D), ASBr (E) and LGR-991 (F) on rice exudates in inducing prehaustorium formation. 20 μM TFBQ 10 μM ASBr (10 μM) and 10 μM LGR-991 were used. Scale bars: 100 μm for A, B, C, 500 μm for D, E, F.

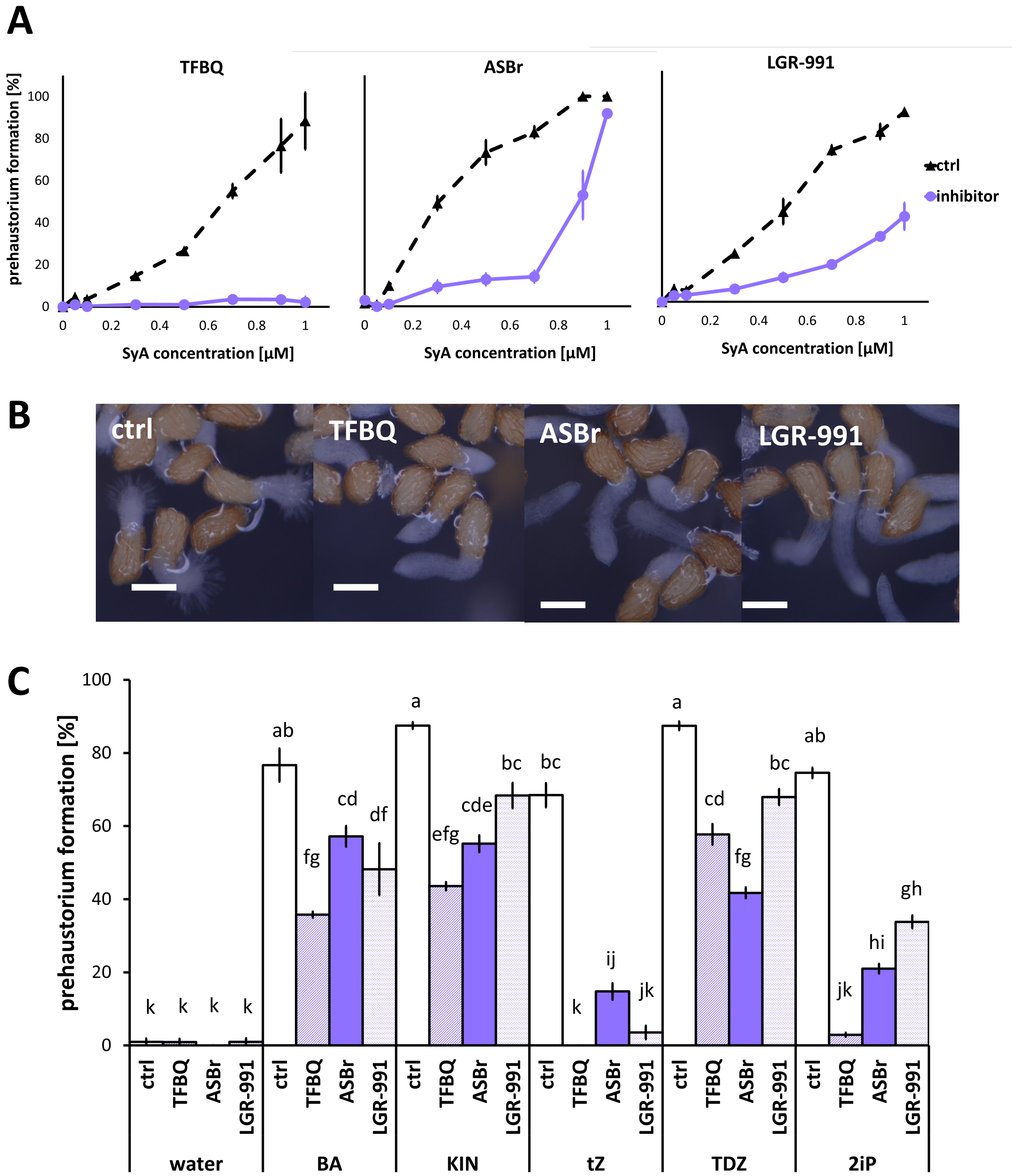


**Figure S3** The effects of chemical inhibitors of HIFs in the presence of SyA or cytokinins on prehaustorium formation in *S. hermonthica*.

A, The effects of TFBQ (20 µM), ASBr (10 µM) and LGR-991 (10 µM) on *S. hermonthica* prehaustorium induction with SyA treatment. B, Images of *S. hermonthica* seedlings treated with SyA in the presence of TFBQ, ASBr or LGR-991. SyA only (ctrl) was used as the control. C, The effects of TFBQ (100 µM), ASBr (10 µM) and LGR-991 (1 µM) on *S. hermonthica* prehaustorium formation induced by BA (300 nM), KIN (500 nM), tZ (25 nM), TDZ (25 nM) or 2iP (100 nM). Data represent the mean ± standard error (n=3). Letters indicate significant differences (Tukey's test, $p<0.05$). Scale bars=100 µm.

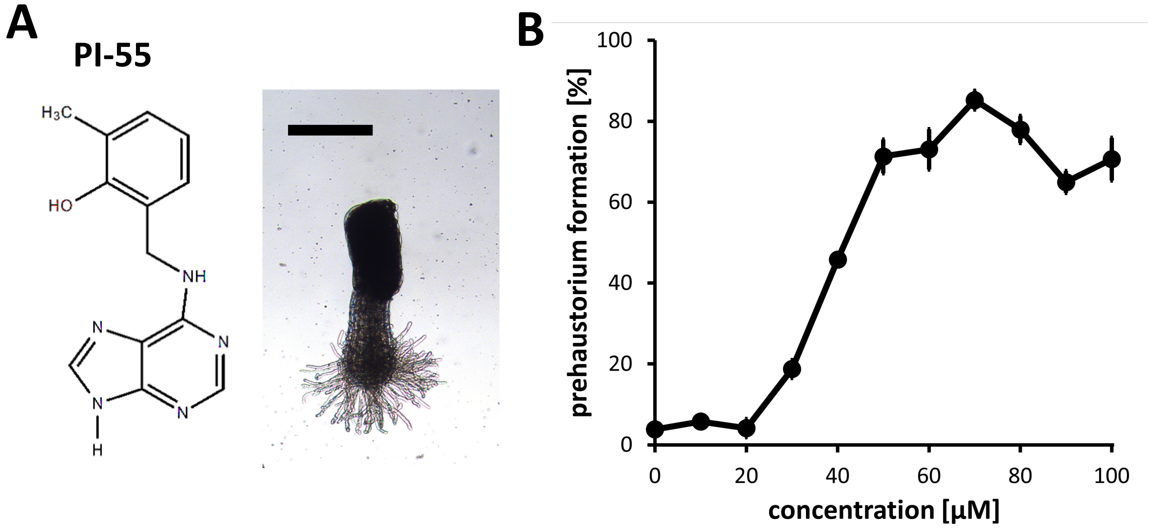


**Figure S4** Cytokinin inhibitors PI-55 and LGR-991 can induce prehaustoria in *S. hermonthica*.
A, The chemical structure of PI-55 (left) and morphology of prehaustoria induced by 10 μM PI-55 in *S. hermonthica* (right). Scale bar = 1 mm. B, LGR-991 induced prehaustoria in *S. hermonthica* at concentrations higher than 20 μM. *S. hermonthica* seedlings were exposed to HIFs and cytokinin inhibitors for 24 h. Data represent the mean ± standard error (n=3).

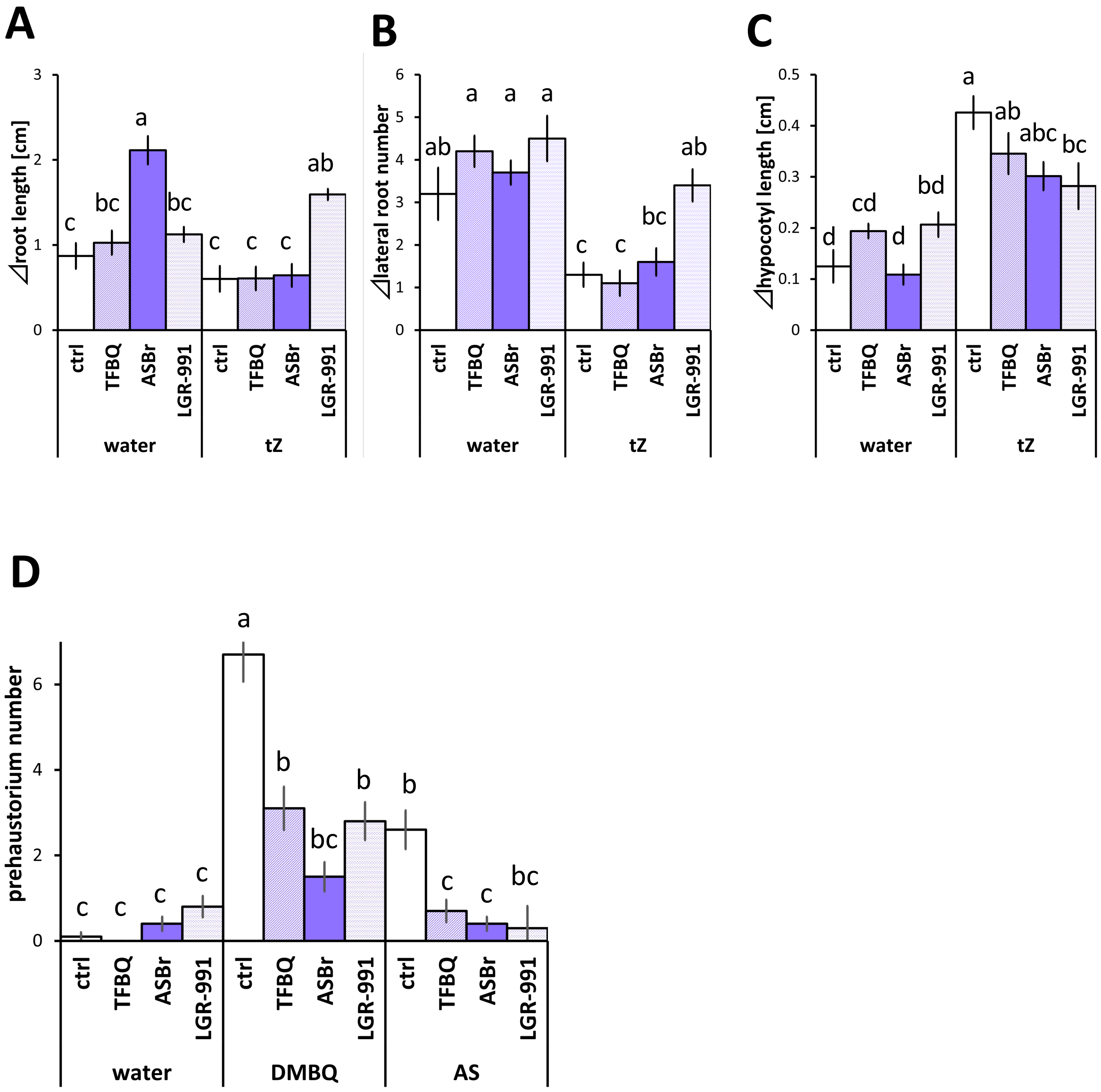


**Figure S5** The effect of inhibitors on prehaustorium induction and cytokinin signaling in *P. japonicum*.
A, The effect of TFBQ (20 μM), ASBr and LGR-991 (10 μM) on *P. japonicum* prehaustorium induction by DMBQ (10 μM) or AS (10 μM). *P. japonicum* seedlings were incubated for 5 days in HIFs and inhibitor solutions. B-C, Effects of TFBQ (20 μM), ASBr and LGR-991 (5 μM) on cytokinin signaling induced by tZ (500 nM) in *P. japonicum*. B, Effects of inhibitors on root elongation suppressed by tZ. C, Effects of inhibitors on lateral root initiation suppressed by tZ. D, Effects of inhibitors on hypocotyl growth promoted by tZ. Data represent the mean ± standard error (n=10). Letters indicate significant differences (Tukey's test, $p<0.05$).

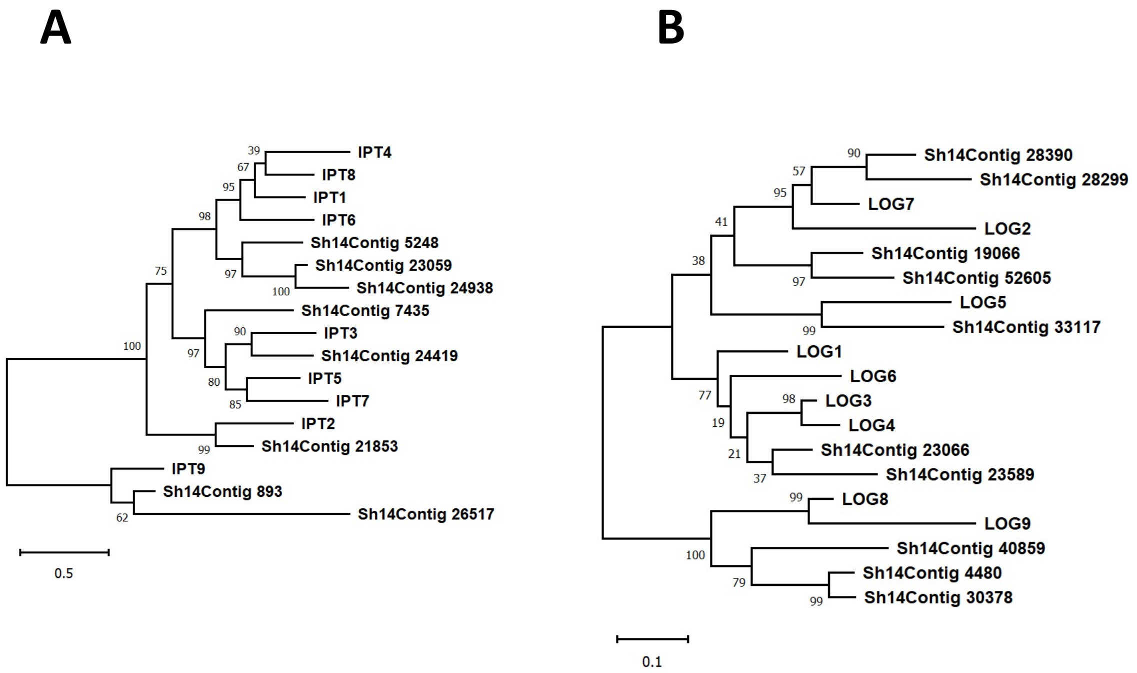


**Figure S6** Phylogenetic trees of cytokinin biosynthesis genes.
A, Phylogenetic tree of IPT genes in *A. thaliana* and *S. hermonthica.* B, Phylogenetic tree of LOG genes in *A. thaliana* and *S. hermonthica.*

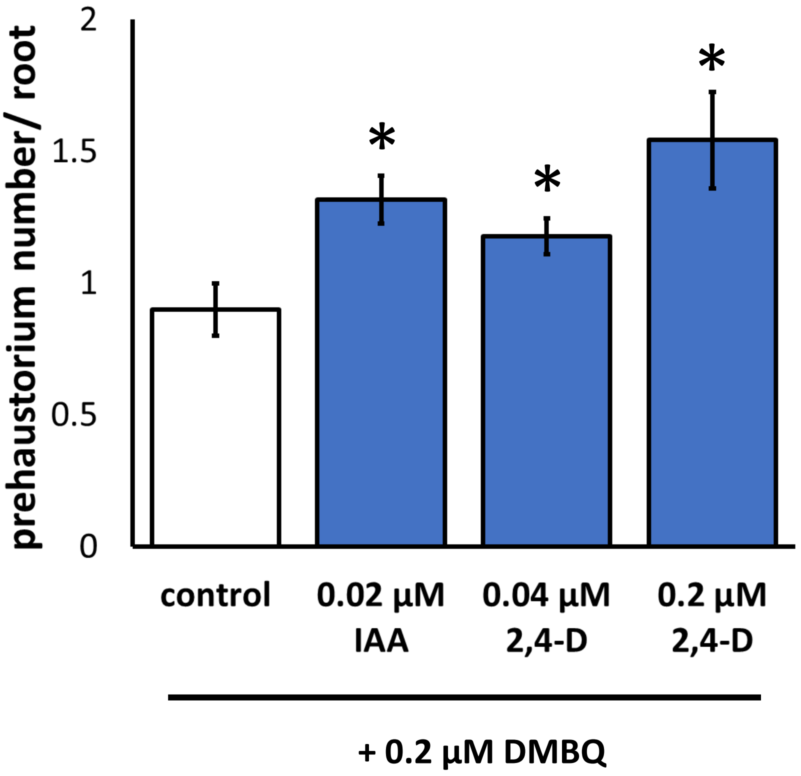


**Figure. S7** The effect of auxin on prehaustorium formation in *P. japonicum.*
A, The effect of auxin on prehaustorium induction by DMBQ. *P. japonicum* seedlings were incubated for 4 days on agar plates containing DMBQ (0.2 µM) and Indole-3-acetic acid (IAA) or 2,4-Dichlorophenoxyacetic acid (2,4-D). Data represent the mean ± standard error (n=22). Asterisks indicate statistically significant differences compared to the control (t-test, $p<0.05$).

**Table S1**. Primer list used in this study

| Primers | Direction | Sequence | Target gene |
|---|---|---|---|
| Sh14Contig_16015 | F | GGATGATGGCCGCTCAATTCAAAG | ShQR2 |
| Sh14Contig_16015 | R | CGCCTTGAGATCCGGTGCTATAAA | ShQR2 |
| Sh14Contig_776 | F | CATGATTCCGGAATGCTTTGGTGA | ShPIRIN |
| Sh14Contig_776 | R | TGGAGGTTAAAGTCATAGCAGGGG | ShPIRIN |
| Sh14Contig_29846 | F | CTTCCACACCTCGACTTTCC | ShYUC3 |
| Sh14Contig_29846 | R | GAACCTTGGGTGGATCTTGA | ShYUC3 |
| Sh14Contig_20216 | F | GAAACGATGTTAACGCGTGCGGAA | ShEXPB1 |
| Sh14Contig_20216 | R | TGGCCCGAGCATATATCCAACGAA | chitinase |
| Sh14Contig_17452 | F | ACCGCGCGGACATTATCGTA | chitinase |
| Sh14Contig_17452 | R | GACGTACGGCCAGATCGTGA | class iv chitinase |
| Sh14Contig_16850 | F | CCTGCCCTCGATTTACTCACTG | class iv chitinase |
| Sh14Contig_13408 | F | ATCCCAGGCATGCAATAGTC | ShRR5 |
| Sh14Contig_13408 | R | AAGGTGACAGCGGTGGATAG | ShRR5 |
| Sh14Contig_22766 | F | CTTTTGAAAGCCTTCGACCTC | ShCKX2 |
| Sh14Contig_22766 | R | CTGGCCCCATTCTCTTCTATC | ShCKX2 |
| Sh14Contig_21853 | F | GTCGTCTGAAAGACCTTGATCC | ShIPT2 |
| Sh14Contig_21853 | R | GGAAGTACACCCAGACGTGAAT | ShIPT2 |
| Sh14Contig_33117 | F | TCTAACAGCTCCTCCAAAGTCC | ShLOG5 |
| Sh14Contig_33117 | R | GGCTAATATGCACCAGAGGAAG | ShLOG5 |
| Sh14Contig_1624 | F | AGCTCCTCTTAATCCCAAGGCCAA | ACTIN (internal control) |
| Sh14Contig_1624 | R | TGACACCATCACCAGAATCGAGCA | ACTIN (internal control) |
